# Fatty acids synergize with bacteriophages against multidrug-resistant pathogens

**DOI:** 10.64898/2026.10.27.759484

**Authors:** Kayla R. Raygoza, Andrew Saghir, Yu Chen, Faith Cribbs, Ritwik Kumar, Liana Hase-Penn, Kyle Kim, Alisha Monsibais, Gina Faraci, Katrine L. Whiteson, Sage J. B. Dunham

**Affiliations:** Department of Molecular Biology and Biochemistry, University of California, Irvine, California, USA

**Keywords:** Bacteriophages, Phage Synergy, Phage Adjuvants, Medium Chain Fatty Acids, Antimicrobial resistance

## Abstract

In the face of rising antimicrobial resistance (AMR), phage therapy presents a promising alternative to antibiotics, but efficacy is limited by biofilms, narrow host range, and the rapid development of phage resistance. Here we present a new concept for phage-based antimicrobials: using small molecules to help phages infect and kill bacteria. Using checkerboard synergy assays, growth curves, and biofilm assays, we evaluate short- and medium-chain fatty acids (SCFAs/MCFAs) with diverse lytic phages against 18 strains of *Stenotrophomonas maltophilia* and extend our analysis to include more than 120 clinical isolates of *Staphylococcus aureus*, *Enterococcus* spp., and *Pseudomonas aeruginosa*. pH-neutral MCFAs (C6–C10) are bactericidal, do not diminish phage viability, and synergized with phages to delay or prevent phage-resistant bacterial rebound over 48 h. MCFAs lowered both attached and free bacterial biomass and were synergistic with envelope-active antibiotics (polymyxin B, ceftazidime, meropenem), supporting a mechanism involving bacterial envelope disruption. Phage-MCFA cooperativity was observed across both Gram-negative and Gram-positive bacteria: across 18 *S. maltophilia* strains and four lytic phages, phage-MCFA combinations were predominantly additive-to-synergistic; in 91 strains of *Enterococcus* spp., pH-neutral C8 produced widespread synergy with phage CAR, and enabled killing of numerous strains that were recalcitrant to phage alone, effectively expanding the functional host range of phage CAR; in 18 strains of *S. aureus*, effects were stoichiometry-dependent, showing both synergy and antagonism depending on the ratio of phage-to-MCFA; in 14 strains of *P. aeruginosa*, interactions were largely additive. Together, this work elevates MCFAs as potential therapeutic accompaniments that stabilize and broaden phage efficacy by addressing several of phage therapy’s limitations at once and highlights the general potential of small molecules as phage adjuvants.

**SIGNIFICANCE:** Bacteriophages (phages) offer a promising therapeutic approach against antimicrobial resistant pathogens, but they are often limited by biofilms, narrow host ranges, and phage resistance. Using multiple *Stenotrophomonas maltophilia* strains and lytic phages, we find that pH-neutral medium-chain fatty acids (MCFAs) enhance phage-mediated killing, reduce biofilm burden, and delay or prevent phage resistance. These benefits generalize beyond *S. maltophilia*: phage-MCFA interactions are broadly synergistic in *Enterococcus* spp., stoichiometry-dependent in *Staphylococcus aureus*, and largely additive in *Pseudomonas aeruginosa*. MCFAs also cooperate with envelope-active antibiotics and are antagonistic with translation inhibitors. Because fatty acids are inexpensive, biocompatible, and straightforward to formulate, our work establishes a practical strategy to create phage-permissive states and improve outcomes of phage, antibiotic, and phage-antibiotic therapies against high-priority infections.

## INTRODUCTION

Antimicrobial resistance (AMR) is one of the biggest challenges facing modern medicine, accounting for approximately 4.71 million associated deaths annually (Naghavi 2024). AMR is a particular threat for people with compromised immune systems and those with cystic fibrosis (CF). CF causes the accumulation of a thick, sticky mucus in the airways; creating an environment that supports persistent infection and disrupts the long-term effectiveness of antibiotics (Perikleous 2023, Dunham 2026). One solution to AMR is found in bacteria’s natural predator: the bacteriophage (McCutcheon & Dennis 2021, Moon 2025). Bacteriophages (or simply, phages) are viruses that exclusively infect bacteria, and are typically active against only a few strains of a single species (Koskella & Meaden 2013, Clokie 2011). This specificity makes phages excellent tools for targeted therapy without the off-target effects of broad-spectrum antibiotics. Phage therapy has a long history of success and phages are generally regarded as safe (Uyttebroek 2022), however their clinical use in most of the world remains limited by challenges such as the development of bacterial resistance and narrow host ranges. To overcome these limitations, researchers typically employ combinatorial approaches, such as multi-phage cocktails and phage-antibiotic co-treatments (Eskenazi 2022, Monsibais 2025, Van Nieuwenhuyse 2022). In the United States, phages are typically used as a compassionate-use therapy of last resort, and are applied in conjunction with the standard of care (usually intravenous antibiotics) (Pirnay 2024).

Inspired by antibiotic adjuvants – which enhance antibiotic activity through a variety of mechanisms shared with phages, such as the inhibition of the bacterial immune response (Dhanda 2023) and the disruption of cell membranes and biofilms (Lin 2023) – we sought to test small molecules as accompaniments to phage therapy. While the use of small molecules to improve phage activity is underexplored, researchers have long understood that small molecules can affect phage biology. Phage biologists have established that simple supplementation with divalent cations such as Mg^2+^ and Ca^2+^ can dramatically enhance adsorption efficiency, genome injection, and overall lytic activity, underscoring how small molecules can potentiate phage infection (Rountree 1955, Ma 2018, Elshazly 2025). Recent work has expanded this principle to deliberate chemical modulation of phage-host interactions. Wong and Balskus demonstrated that diverse microbiome-derived metabolites – including aromatic acids, quorum-sensing molecules, and redox-active compounds – can tune phage infection outcomes by altering bacterial stress responses, cell surface architecture, or prophage regulatory circuits (Wong 2025). Their analysis highlights that small molecules can both enhance and suppress phage infection depending on context, and suggests that the bacterial metabolome may offer a library of small molecules for improving phage efficacy. Complementing this, Zang et al. showed that specific metabolites can inhibit bacterial innate immune systems (notably CBASS), thereby increasing bacterial susceptibility to phage predation (Zang 2026), and Barrio-Pujante et al. demonstrated that the plant-derived auxin indole-3-acetic-acid modulates quorum sensing and suppresses anti-phage defense mechanisms in *Klebsiella pneumoniae* (Barrio-Pujante 2026). Collectively this growing body of work supports a broader hypothesis that exogenous small molecules can act as phage-enhancing agents that weaken bacterial defenses, alter membrane properties, or reshape the physiological environment in ways that favor productive infection.

Widely used as antimicrobial agents in agriculture, food safety, and consumer products, short and medium chain fatty acids (SCFAs and MCFAs) can reduce bacterial load by disrupting membrane integrity, collapsing membrane potential and energy metabolism, and inhibiting biofilm formation (Zhang 2019, Desbois and Smith 2009). Their effects extend beyond simple acidification: recent work demonstrates that commensal-derived SCFAs profoundly remodel *S. aureus* membrane lipid composition, lowering the branched- to straight-chain FA ratio, depolarizing the membrane, suppressing *agr* quorum signaling, and potentiating the activity of membrane-targeting antimicrobials and host peptides across diverse clinical isolates (Fletcher 2026). MCFAs like caprylic acid are generally recognized as safe, and have an established history of human use in nutritional supplements, topical antimicrobials, and pharmaceutical formulations, (FDA 1997, Li 2019). Although SCFAs such as acetate, propionate, and butyrate have not been detected in healthy lungs, they are routinely observed at low millimolar concentrations in sputum from clinically stable people with CF (Ghorbani 2015), suggesting that airway-localized microbial communities naturally generate FAs capable of reshaping pathogen physiology.

Given their natural occurrence, relative safety, and capacity to modulate bacterial defenses, FAs offer a promising yet largely unexplored avenue for enhancing phage efficacy (Kotlyarov 2022, Verma 2024). By weakening bacterial membranes, reducing biofilm barriers, and inducing metabolic or regulatory stress, we expect FAs to create cellular states more permissive to phage adsorption and genome injection. In this study, we evaluate whether FAs improve phage activity against both Gram-negative (*Stenotrophomonas maltophilia* (STM) and *Pseudomonas aeruginosa*) and Gram-positive (*Staphylococcus aureus* and *Enterococcus* spp.) pathogens. Long considered the canonical pathogens of CF, *P. aeruginosa* and *S. aureus* infect approximately 23% and 60% of people with CF respectively (CFF 2024 Patient Registry Highlights, CFF 2023 Patient Registry Annual Data Report) while STM only infects about ∼5% (CFF 2024 Patient Registry Highlights, Terlizzi 2023). Despite its relatively low prevalence, STM’s intrinsic resistance mechanisms – including multidrug efflux pumps, β-lactamases, and a high propensity for biofilm formation – severely limit treatment options and underscore the need for novel therapeutic approaches (Waters 2011, Pompilio 2011, Ryan 2009). *Enterococcus* spp. are increasingly responsible for non-respiratory infections in people with CF (and others), especially urinary tract infections (Reasoner 2022).

To guide our investigation, we focused on three central questions: (1) Can fatty acids interact with bacteriophages to enhance antibacterial activity? (2) How widespread and robust are these effects across strains, species, growth conditions, and phage types – and which factors determine when synergy versus additivity or antagonism emerges? (3) Which broad biophysical changes induced by fatty acids (for example, membrane perturbation or matrix remodeling) govern how phages engage and act on bacterial populations? These questions provide the framework for a systematic assessment of FAs as both antimicrobial agents and potentiators of phage and antibiotic activity across diverse AMR pathogens.

## RESULTS

Bacterial cultures and phage-FA combinations were incubated statically at 37℃ inside humidified, sealed containers to reduce evaporation and create oxygen-limited conditions that better approximate the viscous, low-oxygen microenvironments characteristic of airway infections (see Materials and Methods). Because unbuffered acids can kill bacteria and impact antimicrobial efficacy through acidification alone (Gallagher 2019), FAs were prepared as pH-neutral mixtures of each acid and its conjugate base, ensuring that any observed effects reflected intrinsic FA activity rather than changes in pH. Our collection of phages, strains, fatty acids and antibiotics used in this study are outlined in **Supplementary Tables 1-3**.

### Medium chain fatty acids are synergistic with phage ANB28, while short chain fatty acids are not

We began by performing a series of checkerboard synergy assays against *S. maltophilia* strain K279a using phage ANB28 and the 10 smallest, unsaturated, straight-chain FAs in pH-neutral conditions (**Figure 1**). Although infectious against K279a, phage ANB28 only inhibits growth at a multiplicity of infection (MOI) of approximately 0.01 or higher (approximate titer of 1 × 10^5^ pfu/mL in **Figure 1**) (Monsibais 2025). This intermediate infectivity presents a suitable phage-host pairing for evaluating the effect of different antimicrobials.

**Figure 1.**
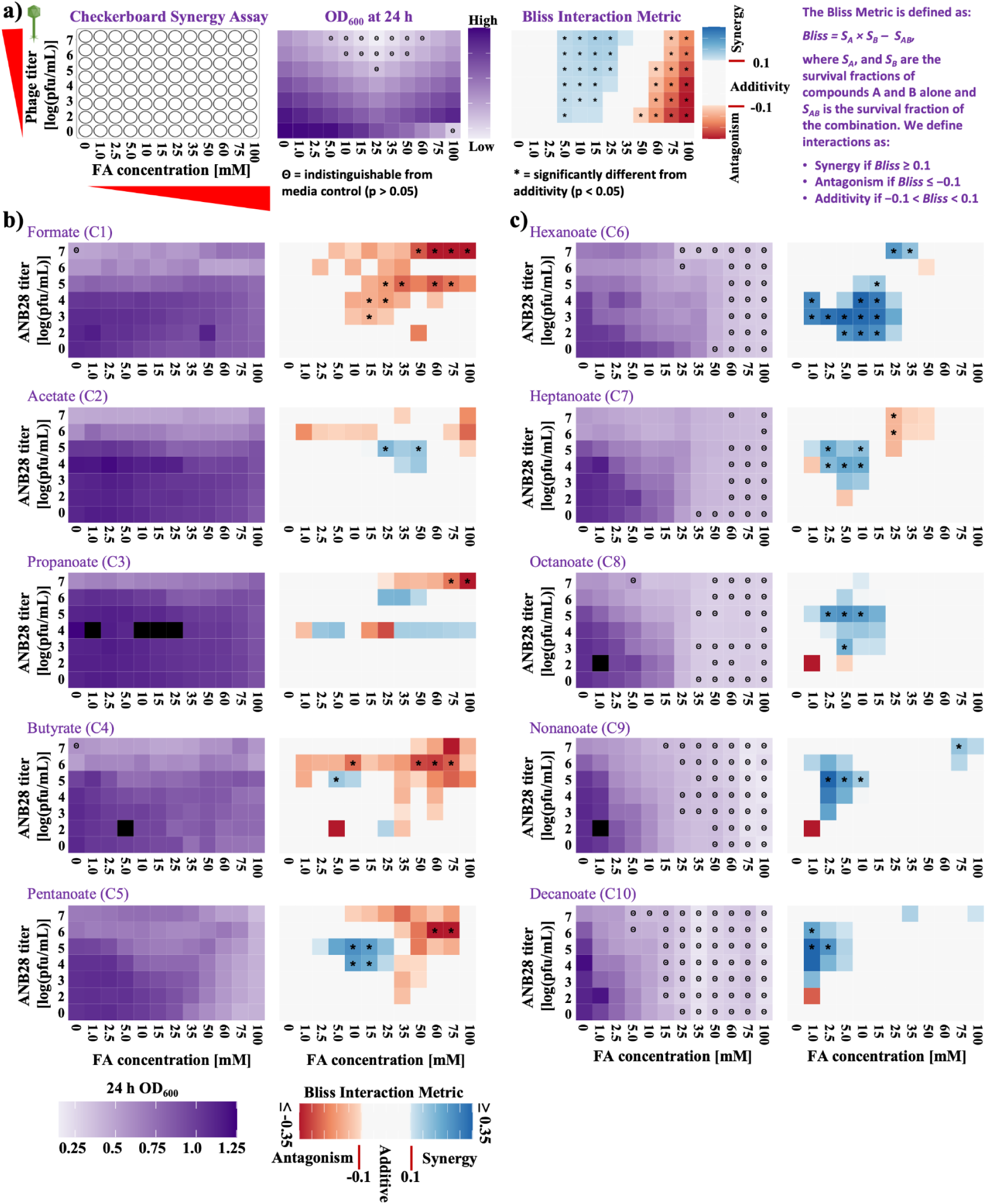
Medium chain fatty acids are synergistic with phage ANB28; short chain fatty acids are not. Checkerboard synergy assays were performed between phage ANB28 and the ten smallest unbranched saturated fatty acids (C1-C10) against *S. maltophilia* K279a. **(a)** Schematic of the assay workflow. **(b)** OD_600_ heatmaps and Bliss Interaction profiles for SCFAs (C1-C5). **(c)** OD_600_ heatmaps and Bliss Interaction profiles for MCFAs (C6-C10). Each plate contains increasing phage titers by row and increasing concentrations of pH-neutral fatty acid/conjugate base by column. After 24 h of growth under oxygen-limiting conditions, OD_600_ values are recorded and Bliss Interaction Metric is calculated for every phage-fatty acid combination. Each assay was performed in triplicate and the average is shown. The highest phage input (1 × 10^7^ PFU/mL) corresponds to an approximate MOI of 1. For the Bliss Interaction Metric, positive values denote synergy, negative values denote antagonism, and values near zero indicate additivity. OD_600_ values that are statistically indistinguishable from media-only controls (p > 0.05, one-sided t-test) are marked with “Θ,” and Bliss Interaction Values significantly different from additivity (p ≤ 0.05 by a one-sample non-equivalence t-test against the interval (−0.1, 0.1)) are marked with “*”.

In **Figure 1**, we show each FA as evaluated through a 24-hour OD_600_ and Bliss Independence heatmaps. Squares on the Bliss Independence map represent the growth response of the bacteria to various stoichiometries of the phage and the FA, with independent (additive) interactions shown as white, synergistic interactions shown as blue, and antagonistic interactions shown as red (**Figure 1a**; see Materials and Methods). In the hypothetical example (**Figure 1a**) we observe additivity (white in the Bliss map) at low concentrations of acid, both additivity and synergy (blue in the Bliss map) at intermediate concentrations, and antagonism (red in the Bliss map) at high concentrations.

Each of the 5 smallest FAs tested (C1-C5; **Figure 1b**) moderately inhibits K279a growth on their own, but interactions between the phage and the SCFAs do not show clear trends. Formate (C1) appears to be antagonistic with phage ANB28, particularly at intermediate concentrations of the phage and FA. Acetate (C2), propanoate (C3), and butyrate (C4) each show both antagonism and synergy at closely related stoichiometries, without any clear trends. Pentanoate (C5) shows a cluster of synergy at around 10 mM and 1 × 10^5^ pfu/mL ANB28 (approximate MOI of 0.01), additivism at lower concentrations, and antagonism at high concentration and titers.

As the FA chain length increases however (**Figure 1c**), the FAs become more inhibitory on their own and display consistent synergistic interactions with the phage. When paired with 1 × 10^5^ pfu/mL ANB28, as little as 2.5 mM of heptanoate (C7) or octanoate (C8) is sufficient to reduce bacterial growth by 50% at 24 hours. Similarly, when used in conjunction with 1 × 10^5^ pfu/mL ANB28, the lowest concentration of decanoate (C10; 1mM) reduced growth by more than 50%. When used without phage, 10 mM C8, 2.5 mM C9, and 2.5 mM C10, was needed to achieve 50% growth reduction (**Figure S1**). Taken together, these results indicate that MCFAs synergize with phage ANB28 to inhibit growth of *S. maltophilia* strain K279a. Further, at stoichiometries where synergy is not observed, the interactions appear to be mostly additive which is also a favorable outcome from a therapeutic standpoint.

### MCFAs inhibit STM growth independent of pH

We compared the inhibitory effects of the same 10 FAs as pH-neutral mixtures of each acid and its conjugate base (as in **Figure 1**) to their activity in their acidic form, by allowing the pH to fluctuate after the buffering capacity of the BHI media was exceeded (**Figure S1**). Using pH neutral solutions of each FA and its conjugate base (**Figure 1** and **Figure S1a**), SCFAs (C1-C4) exhibited only moderate growth inhibition, even at concentrations as high as 100 mM. Meanwhile, MCFAs retained potent antimicrobial activity: as little as 2.5 mM of C9 or C10 caused greater than 50% growth reduction, and both compounds showed measurable effects even at 1 mM.

In their acidic form (**Figure S1b**), all tested FAs passed the 50% growth reduction threshold at concentrations ranging from 15-50 mM. For SCFAs with 5 carbons or less, 95% growth inhibition occurred at concentrations that exceeded the buffering capacity of the media, causing the pH to fall below 6 (**Figure S1b-c**). C6 was also more effective in the acidic form, with 25 mM needed to achieve 95% growth inhibition (compared to 65 mM under pH neutral conditions), while C7 was similar under both pH conditions (65 mM to achieve full growth inhibition). In contrast, C8 and C9 exhibited superior growth inhibition under pH neutral conditions (35 mM and 25 mM versus >100 mM). We were not able to reliably solubilize C10 in its acidic form, but the pH-neutral combination of the acid and its conjugate base achieved full growth inhibition at 25 mM and 50% growth inhibition at 2.5 mM. These results indicate that while SCFA-mediated inhibition is largely pH-dependent, MCFAs inhibit STM growth through a mechanism that is (at least in part) independent of pH.

### MCFAs delay or prevent the emergence of phage-resistant STM across multiple strains, phages, and media compositions

Given that the straight-chain MCFAs tested above exhibited similar antimicrobial and phage-enhancing behavior, we used pH-neutral C8 (which we will refer to as OCT) as a representative compound to study how MCFAs influence phage suppression during long-term growth. We monitored 48-h growth dynamics for two *S. maltophilia* strains (K279a and B28B), each paired with an active phage (ANB28 or SBP2Φ2, respectively), in both Brain Heart Infusion (BHI) and Synthetic CF Sputum Media (SCFM1).

Examining strain K279a first (**Figure 2a-b**), both ANB28 and OCT activity differed markedly between media types. At an MOI of 10, the phage effectively suppressed growth in BHI up to 24 h followed by a strong rebound (**Figure 2a**), and was largely ineffective in SCFM1 (**Figure 2b**). In both media types OCT alone reduced growth in a concentration-dependent manner, and was much more effective in SCFM1 than BHI: 10 mM resulted in ∼90% growth suppression at 48h in SCFM1, whereas this same concentration resulted in ∼20% 48h growth suppression in BHI. Combining ANB28 with OCT substantially altered long-term growth kinetics in both BHI and SCFM1, with 10 mM OCT and ANB28 sufficient to hold cultures at or near baseline OD for the duration of the experiment in both media types. At lower MOIs (∼1 and 0.1) phage ANB28 was largely inactive on its own, but strongly synergistic with the MCFA (**Figure S2**). These results indicate that OCT slows or prevents the expansion of ANB28-resistant subpopulations in K279a, including in CF-mimicking media.

**Figure 2.**
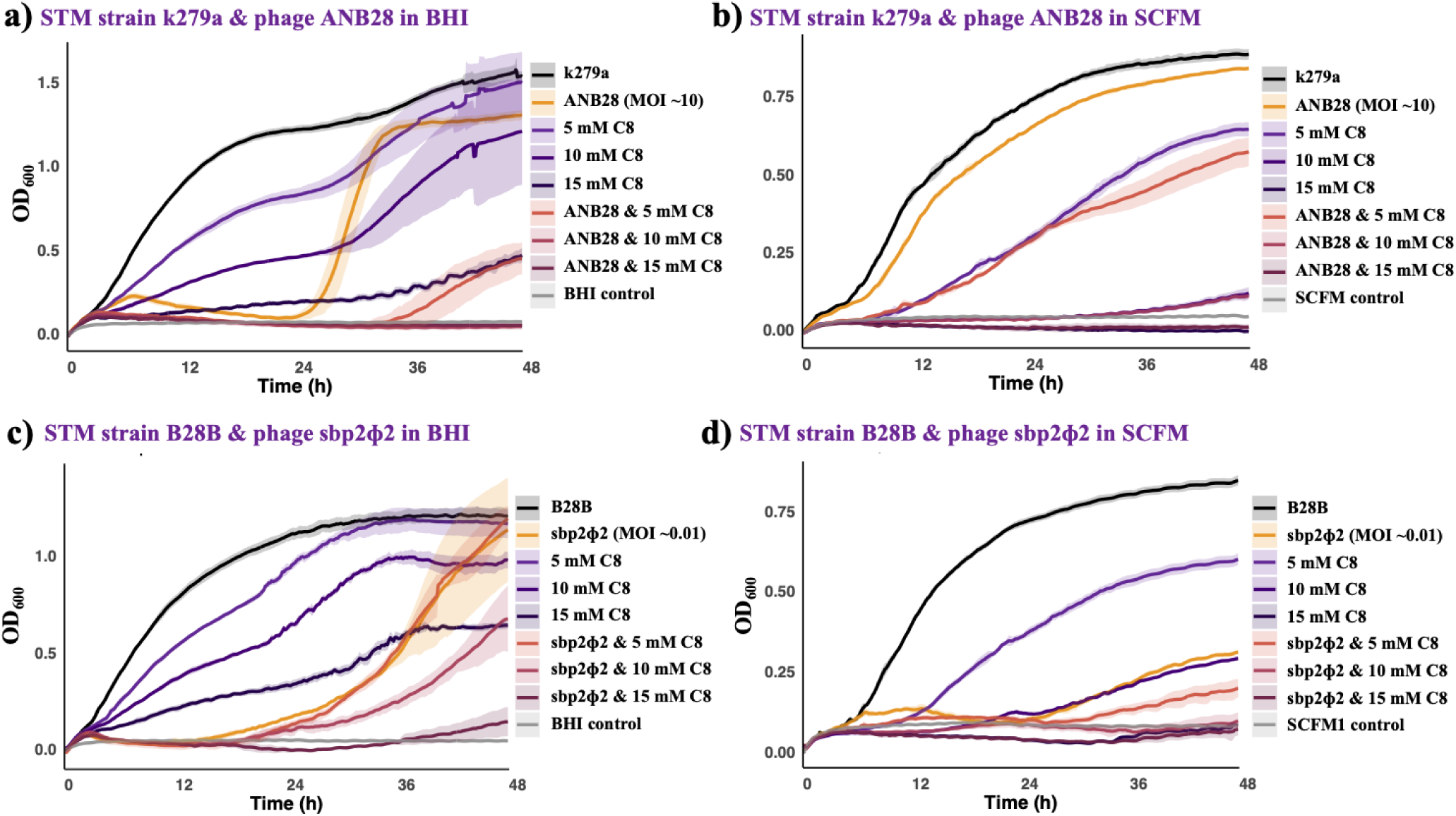
MCFAs prevent the development of phage resistance and are active in SCFM. 48 h growth curves for STM strain K279a and phage ANB28 in (**a**) BHI and (**b)** SCFM, with exposure to varying amounts of OCT. Analogous curves for STM strain B28B and phage SBP2ɸ2 in (**c**) BHI and (**d**) SCFM. Shown are the averages (solid lines) and standard deviations (shading) for 5 replicates. Phages were added at an approximate titer of 1 × 10^8^ pfu/mL for ANB28 and 1 × 10^5^ pfu/mL for SBP2ɸ2.

For STM strain B28B, a similar pattern emerged with phage SBP2Φ2 (**Figure 2c–d**). Consistent with our prior work (Monsibias 2025), SBP2Φ2 exhibits rapid adsorption and a long latent period in BHI, leading to early host suppression followed by a characteristic rebound where resistant mutants expand and reach culture densities similar to the untreated control by 48 hours. In BHI (**Figure 2c**), OCT alone produced a concentration-dependent reduction in growth and endpoint OD, with 5 mM allowing near-complete regrowth by ∼36 h, while 10 mM and 15 mM partially reduced growth to ∼75% and ∼50% of the control, respectively. In combination with the phage, however, OCT dramatically increased the longevity of SBP2Φ2 activity. At 5 mM, little improvement over phage alone was observed, whereas 10 mM delayed rebound and reduced final culture density by roughly half compared to the untreated control. At 15 mM, bacterial growth was strongly suppressed, with only limited rebound emerging after ∼36 h. Compared to the K279a-ANB28 pairing in BHI, where combination treatment with just 5mM drastically reduced final OD, the synergy found in the B28B-SBP2Φ2 pairing is more limited, consistent with differences in baseline phage efficacy and host response.

In SCFM1, however, the interaction between SBP2Φ2 and OCT was markedly enhanced (**Figure 2d**). SBP2Φ2 alone remained active, suppressing growth more effectively than in BHI and limiting rebound to ∼30% of the untreated control. Once again, OCT displayed increased potency in SCFM1, with 10 mM reducing growth to ∼30% of control levels and 15 mM nearly abolishing detectable growth. In combination, 5 mM OCT greatly enhanced phage-mediated suppression relative to either treatment alone, while 10 and 15 mM OCT either abolished rebound entirely or reduced the final growth to a small fraction of the control. Notably, this contrasts with the K279a-ANB28 system, where phage activity was diminished in SCFM1, indicating that media-dependent effects on phage efficacy are phage-host specific.

Checkerboard synergy assays were performed in both BHI and SCFM against the two STM strain-phage combos used in the growth curves of **Figure 2**, as well as a third strain (SM49), which exhibits intermediate susceptibility to phage SBP2Φ2 (**Figure S3**). As expected, OCT alone inhibited growth of all three bacterial strains, with more pronounced inhibition in SCFM than in BHI. When OCT was combined with phages, bacterial inhibition was generally enhanced relative to either treatment alone. Bliss interaction analyses revealed that most OCT-phage combinations in SCFM clustered near zero, consistent with additive (independent) effects rather than strong synergy or antagonism (**Figure S3**, right panels). In BHI, the additive-to-synergistic interaction pattern was observed across all three STM strain-phage pairs. Importantly, widespread antagonism was not detected in either media composition, suggesting that OCT does not impair phage activity.

Across both of these assays and all strains, phages, and media, the dominant observation is clear: MCFAs consistently increase the long-term stability of phage-mediated suppression by delaying or preventing the emergence of phage-resistant populations. These effects are dose-dependent and more pronounced in SCFM1, suggesting that MCFAs act not only as direct antimicrobials but also as potent stabilizers of phage efficacy under physiologically relevant conditions.

### MCFAs are bactericidal but do not damage phages

We evaluated if the pH-neutral MCFAs were bacteriostatic or bactericidal. Using OCT as a representative MCFA, *S. maltophilia* strain K279a was incubated for 24 hours with varying concentrations of OCT, washed to remove residual OCT, and plated for colony forming unit (CFU) enumeration. As can be seen in **Figure S4a**, there is a sharp drop in both OD_600_ and the number of viable cells present between 10 and 15 mM OCT, and by 50 mM the bacteria are unculturable. The loss in cell viability following OCT removal leads us to conclude that the MCFA is permanently killing the bacteria rather than temporarily inhibiting their growth.

To assess the viability of phage ANB28 in the presence of OCT, we incubated the phage in SM buffer with different concentrations of OCT for two hours at 4℃ (common phage storage conditions), followed by a soft-agar overlay assay with *S. maltophilia* K279a (**Figure S4b**). The resulting phage titers remain within an order of magnitude, regardless of the concentration of OCT. This suggests that even the highest concentration of pH-neutral OCT (100 mM) does not substantially impact phage viability.

### Synergistic effects of MCFAs are derived – at least in part – from biofilm disruption

To evaluate if the observed MCFA-phage synergy was derived from a reduction in biofilm formation, tubes containing bacteria only, bacteria and phage, bacteria and OCT, and bacteria, phage and OCT (combo treatment), were incubated for 48 hours. OD_600_ readings were taken at 24 and 48 h, and the culture was decanted, leaving only attached biofilm which was stained with crystal violet for a measure of attached biofilm. The decanted culture (containing live and dead cells, extracellular polymeric substances, and other biological material from the liquid phase, tube bottom, and liquid–air interface) was added to another tube, homogenized, and the OD_600_ was measured for an approximate measure of total, unattached biomass.

Across all time points and biomass measurements, OCT consistently and significantly reduced the bacterial and biofilm loads. At 24 h, phage ANB28, 10 mM OCT, and the combination treatment all significantly reduced bacterial growth relative to the untreated control. No significant additional reduction was observed in the combination, consistent with a ceiling effect rather than synergy (**Figure 3a**). By 48 hours, the untreated culture maintained its substantially greater growth than the other treatments as phage-only, OCT-only, and combination conditions showed similarly low optical densities (**Figure 3b**). However, when total free biomass was measured at 48 h, the phage-only condition showed a significantly higher OD_600_ relative to all other treatments, indicating the accumulation of biomass not captured in standard OD_600_ measurements (**Figure 3c**). In contrast, OCT-treated and combination cultures had lower biomasses, consistent with minimal accumulation of biomass at the bottom or top of the tubes. Notably, the phage treated and OCT treated conditions showed similar 48 hour OD_600_ values (**Figure 3b**) but the phage treated cultures show comparably high levels of free biomass, suggesting that the low biomass in the OCT condition results from something beyond a decline in cell abundance alone.

**Figure 3.**
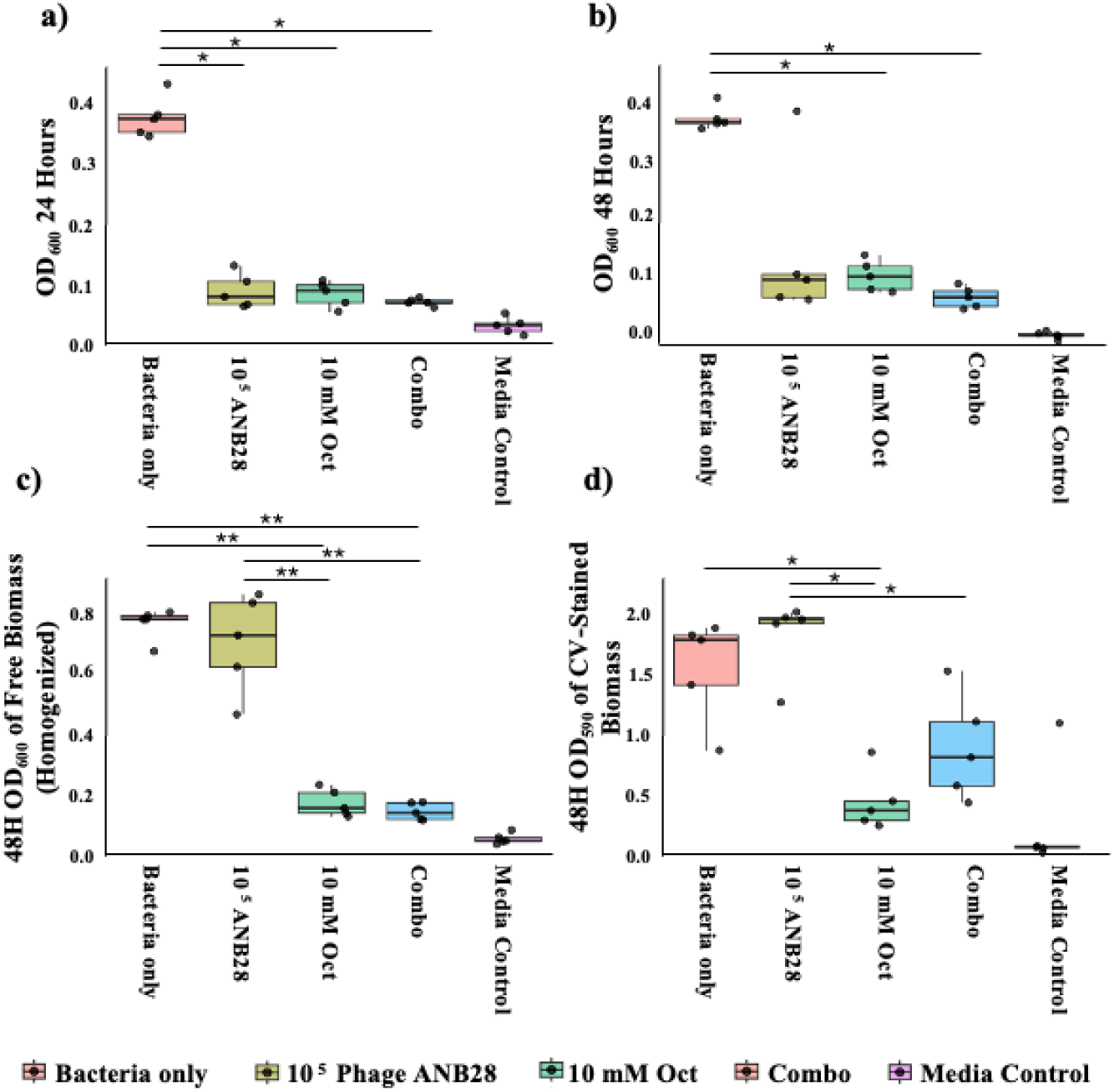
MCFAs inhibit biofilm formation in K279a. **(a)** OD_600_ at 24 hours for 5 replicates of 5 conditions: Bacteria only, 10^5^ phage ANB28, 10 mM OCT, a combination of phage and OCT, and a media control. (**b)** OD_600_ at 48 hours under the same conditions. **(c)** OD_600_ resuspended cultures at 48 hours, capturing total biomass (mixture of living and dead cells, biofilm aggregates on the liquid-air interface and settled biomass). (**d)** OD_590_ of the quantification of surface-adhered biofilm by crystal violet staining followed by acetic acid solubilization. Boxplots show the median and interquartile range (IQR), with whiskers extending to min and max points within 1.5x the IQR. Individual data points are overlaid as jittered points representing 5 replicates. Statistical significance between conditions was assessed using pairwise Wilcoxon rank-sum tests with Benjamini–Hochberg (FDR) correction; *p* < 0.05. * and ** represent *p* < 0.05 and *p* < 0.01, respectively.

Crystal violet staining of attached biofilms further supported these findings: the untreated control and phage-treated cultures formed high levels of attached biofilm, while OCT significantly reduced biofilm formation – both when used alone and in combination with phage (**Figure 3d**). Taken together, these data show that the synergistic effects of MCFAs are derived (at least in part) from a reduction in biofilm formation, which presumably allows easier access to the cell surface by phage ANB28.

### MCFAs work cooperatively with a diverse set of phages to inhibit many STM strains, in both BHI and SCFM

Building on the results of **Figure 1** we evaluated the universality of the antimicrobial and phage enhancement by pH-neutral MCFAs. We tested 5 mM OCT alone and in combination with 4 lytic phages (ANB28, SPB2Φ2, KB824, and ESCSMPI) against 18 strains of STM in both BHI and SCFM1 (**Figure 4**). These assays were performed in 96-well plates, with at least three replicates per condition. Each subfigure shows the average 24 hour OD_600_ value for each bacterial strain and with the addition of phage alone, OCT alone, or both phage and OCT. Average Bliss synergy values are shown as heatmaps and the % growth reduction values for the combined condition are shown as box plots.

**Figure 4.**
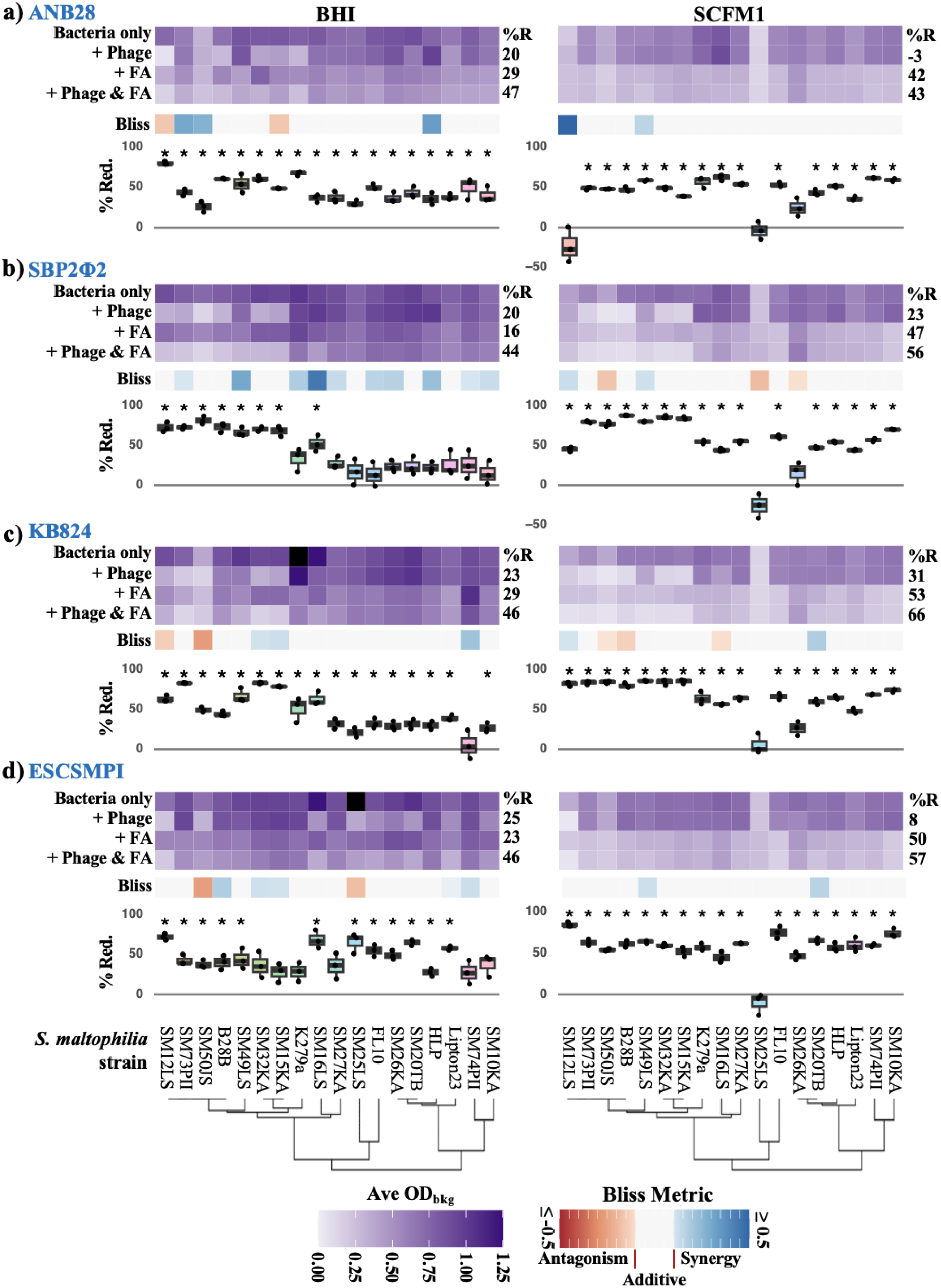
MCFAs work cooperatively with phages to inhibit STM growth across many strains, phages, and media compositions. pH neutral OCT and phage inhibition of 18 strains of *S. maltophilia* in BHI (left) and SCFM (right). Shown are 24 hr OD_600_, Bliss Interaction Metric, and average % growth reduction across all strains from 5 mM pH neutral OCT and four phages: (**a**) ANB28; (**b**) SBP2Φ2, (**c**) KB824, and (**d**) ESCSMPI. The *S. maltophilia* strains are arranged in order of phylogenetic relatedness. Phage was added at 1 × 10^5^ pfu/mL in each case (approximate MOI of 0.01). Positive Bliss values indicate synergy, negative values indicate antagonism, and values near zero indicate additive (independent) interactions. After FDR correction, no Bliss values significantly deviated from additivity (FDR-corrected p ≤ 0.05 by a one-sample non-equivalence t-test against the interval (−0.1, 0.1)). For the % reduction plot the “*” signifies a growth reduction value for the combined condition that was significantly different than 0 (FDR-corrected p-value < 0.05). Data for each experiment is shown as the average of 3 replicates.

In BHI (**Figure 4**, left column), ANB28, KB824, and ESCSMPI inhibited growth in about half of the strains, while SBP2Φ2 was effective against only a third of the strains. KB824 and SBP2Φ2 retained activity in both media types, while ANB28 and ESCSMPI lost efficacy in SCFM1, highlighting strain- and media-specific dependencies. OCT alone consistently suppressed growth across most strains in both media types, with higher average growth reductions observed in SCFM1 (42–53%) compared to BHI (16–29%).

Across nearly all conditions and strains, combining 5 mM OCT with phage treatment yielded additive effects, particularly in strains with intermediate phage susceptibility. In BHI, average growth reduction from combination treatments consistently exceeded that of either treatment alone – for example, ANB28 + OCT caused 47% reduction (vs. 20% and 29% individually), while similar enhancements were seen with SBP2Φ2 (44% vs. 20% and 16%), KB824 (46% vs. 23% and 29%), and ESCSMPI (46% vs. 25% and 23%). In SCFM1, where ANB28 and ESCSMPI were largely inactive on their own, OCT still maintained antimicrobial effects, and the combination treatment improved outcomes in most cases. Antagonistic interactions fell below the threshold of significance and were rare. They primarily occurred when phages already caused near-complete suppression or where bacterial growth was minimal, suggesting they were an artifact of how the Bliss Metric is calculated, rather than a true biological effect. These patterns mirror the cooperative MCFA–phage behavior observed elsewhere in this manuscript, but here extend across four phages and two media compositions. Taken together, these findings suggest that MCFAs not only contribute their own broad-spectrum antimicrobial activity but also reliably improve the efficacy of diverse phages across strains and environments, supporting its potential as a universal phage enhancer against STM.

### MCFAs enhance phage-mediated growth suppression across diverse bacterial genera

Inspired by the broad-strain activity of pH-neutral OCT against *S. maltophilia*, we extended our investigation to test whether MCFAs could similarly enhance phage efficacy against other common pathogens, including both Gram-positive (*Staphylococcus aureus*, *Enterococcus faecalis*, *Enterococcus faecium*; **Figure 5**) and another gram-negative organism (*Pseudomonas aeruginosa*, **Figure S5**). For each species, we first used the multi-strain assay layout (as in **Figure 4**) followed by checkerboard synergy assays for select strains.

**Figure 5.**
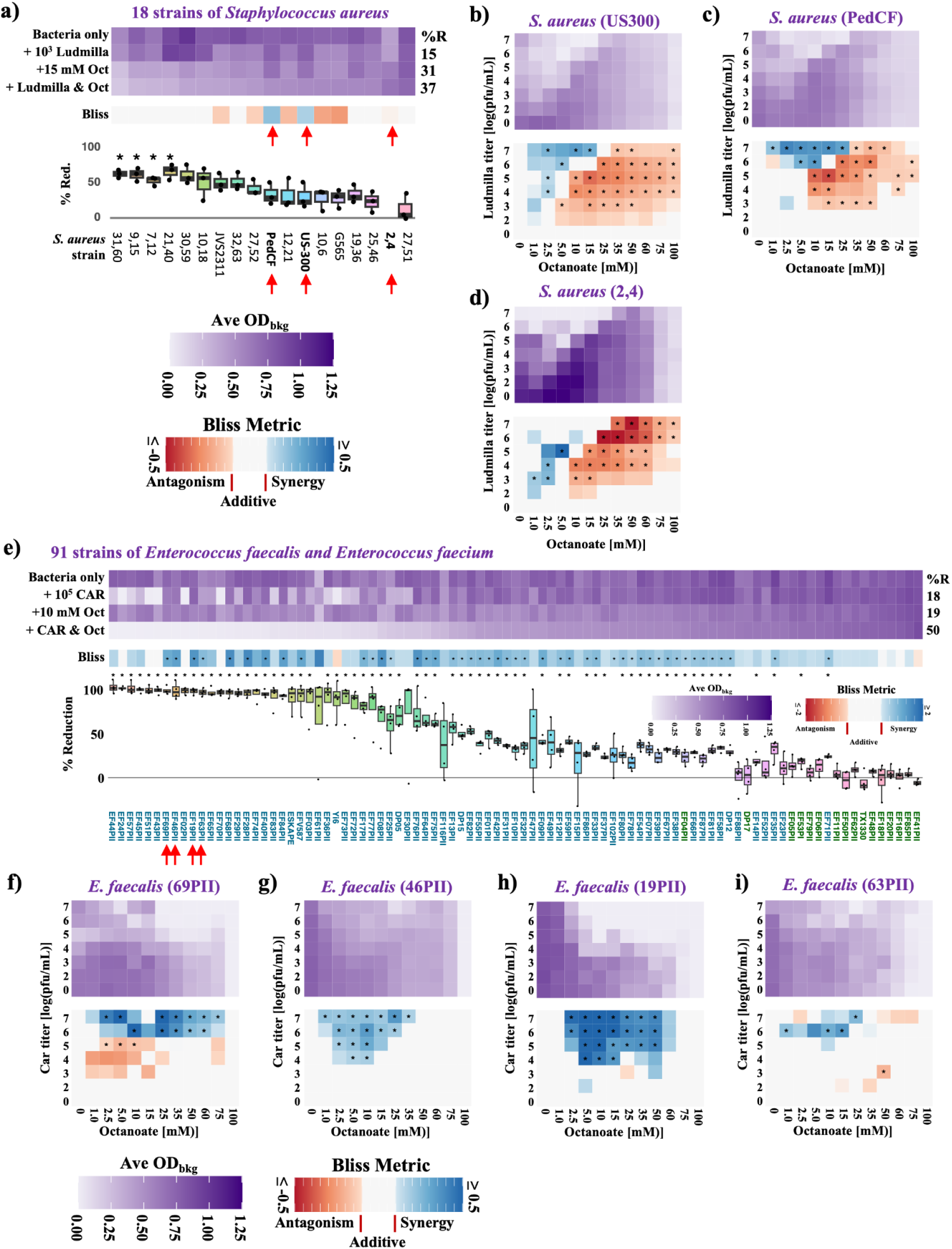
MCFAs exhibit species- and stoichiometry-dependent interactions with bacteriophages in Gram-positive pathogens. **(a)** Multi-strain assay showing 24 h OD_600_ measurements for 18 *Staphylococcus aureus* strains treated with phage Ludmila (10^5^ PFU/mL), OCT (15 mM), or the combination. (**b–d**) Checkerboard synergy assays for three representative *S. aureus* strains (PedCF; USA300; and 2,4), showing OD_600_ heatmaps (top) and corresponding Bliss Interaction Metrics (bottom). (**e**) Multi-strain assay for 91 *Enterococcus* strains (*E. faecalis* and *E. faecium*), treated with *E. faecalis* phage CAR (10^5^ PFU/mL), OCT (10 mM), or their combination. *E. faecium* strains are indicated by green text. (**f–i**) Checkerboard synergy assays for four representative *E. faecalis* strains (69PII, 46PII, 19PII, and 63PII). Positive Bliss values indicate synergy, negative values indicate antagonism, and values near zero indicate additive (independent) interactions. The “*” in the Bliss maps indicate strains with significant positive or negative interactions [p ≤ 0.05 by a one-sample non-equivalence t-test against the interval (−0.1, 0.1), with FDR-correction in **a** and **e**, but not for the checkerboard synergy assays], and for the % reduction plot the “*” signifies a growth reduction value for the combined condition that was significantly different than 0 (FDR-corrected p-value < 0.05). Data for each experiment is shown as the average of 3 replicates, except for panel **f**, which is from 4 replicates.

Across 18 *S. aureus* strains, treatment with the lytic phage Ludmila resulted in measurable growth inhibition in approximately half of the strains tested, corresponding to an average reduction in OD_600_ of ∼15% across the full panel (**Figure 5a**). Consistent with prior observations in *S. maltophilia* (**Figure 4**), OCT exposure alone broadly inhibited *S. aureus* growth, with nearly all strains exhibiting reduced growth at 24 h (average reduction of ∼31%). The combined treatment of OCT and Ludmila produced a modest additional reduction in growth (average ∼37%), suggesting predominantly additive or slightly antagonistic effects at the fixed stoichiometries tested in the multi-strain format.

To more fully resolve interaction dynamics, we performed checkerboard synergy assays on four representative *S. aureus* strains: two clinical isolates (PedCF; and 2,4) and the commonly used laboratory strain USA300 (**Figure 5b–d**). Across all strains tested, the interaction between OCT and Ludmila was strongly dependent on both OCT concentration and phage titer. At lower OCT concentrations, combinations frequently exhibited positive Bliss interaction scores, indicative of synergy, whereas higher OCT concentrations shifted interactions toward antagonism, particularly at lower phage titers. For example, in *S. aureus* strain US300, OCT concentrations of approximately 1 - 15 mM combined with phage titers of 10^6^ - 10^7^ PFU/mL produced strong synergistic interactions, with growth inhibition exceeding that predicted by independent action (**Figure 5b**). In contrast, higher OCT concentrations (10 - 50 mM) combined with lower phage titers (10^2^ - 10^5^ PFU/mL) resulted in marked antagonism. Similar stoichiometry-dependent transitions between synergy and antagonism were observed across all *S. aureus* strains tested, indicating that interactions are highly sensitive to dosing ratios in this species.

We applied the same experimental framework to a larger panel of *Enterococcus* spp. strains, including 91 strains of *E. faecalis* and *E. faecium* (**Figure 5e**). *E. faecalis* phage CAR exhibited activity towards ∼30 of the *E. faecalis* strains in our collection, and none of the *E. faecium* strains. OCT alone (10 mM) produced a consistent reduction in growth across the majority of strains (average of 19%), similar to that observed in *S. aureus* and *S. maltophilia*. Strikingly, the combination of OCT and phage CAR resulted in synergistic growth inhibition across almost every *E. faecalis* strain. In many instances (e.g., EF46 and EF19PII) it appears that OCT helped phage CAR to infect otherwise recalcitrant strains. In contrast, all the *E. faecium* strains showed broadly additive inhibition by CAR and OCT, suggesting that OCT did not expand the host range of CAR beyond its primary host species. Notably no significant antagonistic interactions were observed between phage CAR and OCT.

Follow-up checkerboard synergy assays against 4 *E. faecalis* clinical isolates that showed no activity with CAR alone and strong synergy with OCT – namely EF69PII, EF46PII, EF19PII, and EF63PII (**Figure 5g–j**) – showed broadly synergistic or additive interactions. In the cases of strains EF69PII and EF63PII (**Figure 5g** and **5j**), the checkerboard assays showed that phage CAR was active when applied alone and enhanced by the presence of OCT (EF69PII also showed some significant antagonism at low phage titers). For strains EF46PII and EF19PII (**Figure 5h–i**) we observed that the phage alone was inactive at the titers tested (up to 10^7^ PFU/mL), but strongly synergistic with OCT at concentrations from 1-50 mM.

Together, these data demonstrate that while MCFA-phage interactions are strongly stoichiometry-dependent in *S. aureus*; *Enterococcus* spp. display more stable and broadly synergistic behavior. This contrast highlights species-specific interaction landscapes and suggests that Gram-positive pathogens differ substantially in how MCFAs modulate phage efficacy.

We also examined whether the cooperative effects extended to *P. aeruginosa*, another common Gram-negative pathogen. Treatment with 10^5^ PFU/mL phage PAPS1 alone resulted in noticeable growth suppression of 8/14 strains, yielding an average growth suppression of 23% (**Figure S5a**). Exposure to 15 mM OCT alone produced an average of 31% growth reduction across the panel, with some strain-level variation in effect. Combined treatment with OCT and phage PAPS1 generally resulted in slightly enhanced growth inhibition relative to either agent alone (43% average growth reduction), and Bliss interaction scores clustered near zero for most strains, indicating predominantly additive interactions. Checkerboard synergy assays performed with three representative strains (26,48; PAO1; and P14) confirmed this pattern, revealing largely additive interaction landscapes across a wide range of OCT concentrations and phage titers (**Figure S5b–d**). Together, these data suggest that in *P. aeruginosa*, MCFAs can be combined with phages without broadly impairing phage activity, but they do not consistently drive strong synergistic effects for the strains, phage, and conditions tested here.

### MCFAs are synergistic with antibiotics that disrupt the cell membrane

Inspired by reports of synergy between some FAs and antibiotics (e.g., Lin 2023), we next sought to investigate interactions between fatty acids and antibiotics of diverse mechanisms of action. We performed checkerboard synergy assays pairing pH-neutral C2, C6, or OCT with six clinically relevant antibiotics against *S. maltophilia* strain K279a (**Figure 6**). In agreement with prior observations that SCFAs display minimal standalone activity, C2 did not meaningfully inhibit growth and produced Bliss interaction values near zero across all antibiotic combinations, indicating no detectable impact on antibiotic efficacy. In contrast, both C6 and OCT generated distinct interaction signatures that depended strongly on the antibiotic class.

**Figure 6.**
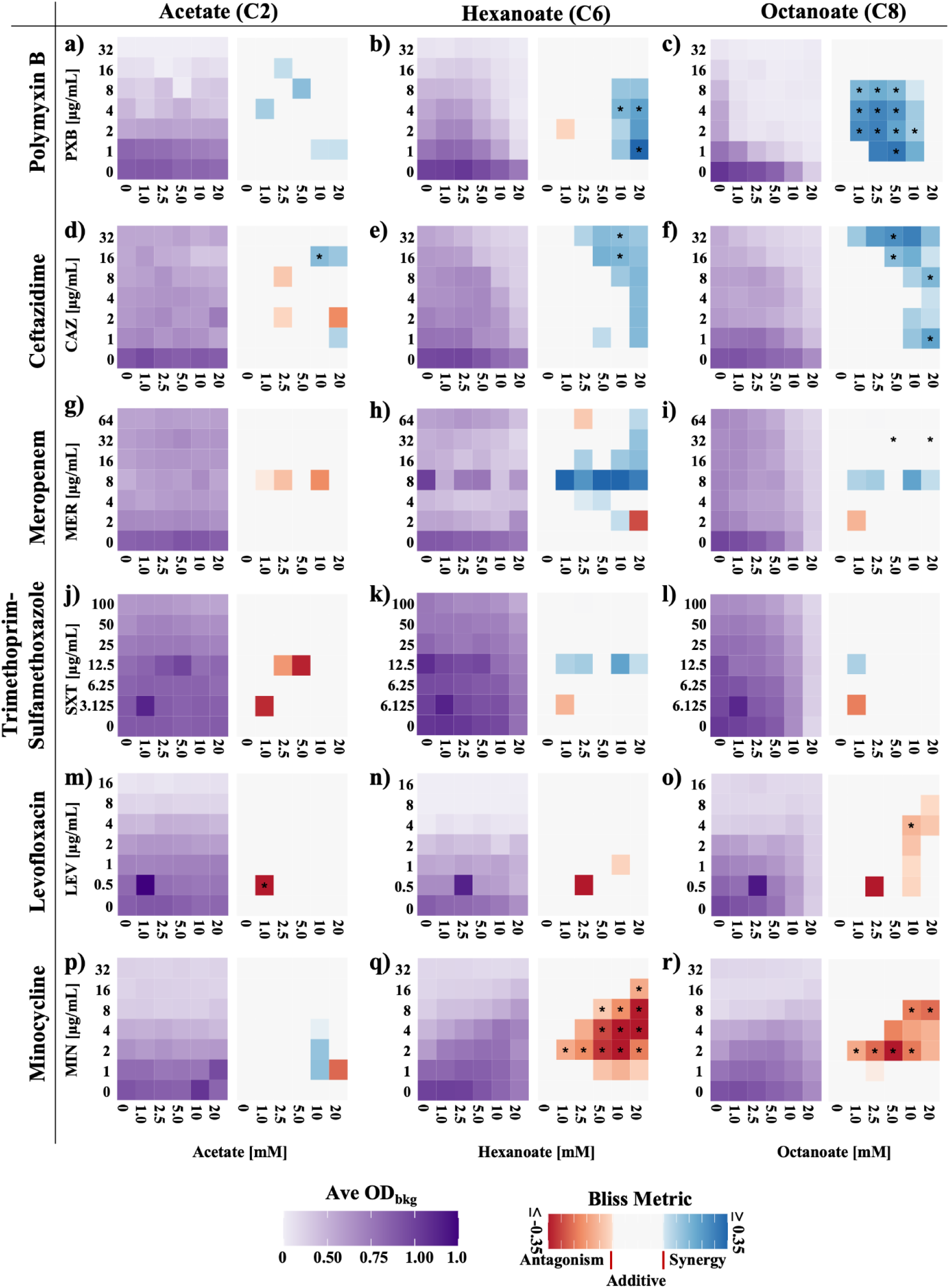
MCFAs are synergistic with antibiotics that inhibit the cell envelope, antagonistic with those that affect intracellular processes. Checkerboard synergy assays between antibiotics and pH-neutral fatty acids on *S. maltophilia* strain K279a. Shown are interactions between three pH-neutral fatty acids (acetate, hexanoate, and octanoate) and six antibiotics: **(a-c)** polymyxin B; **(d-f)** ceftazidime; **(g-i)** meropenem; **(j-l)** trimethoprim-sulfamethoxazole; (**m-o**) levofloxacin; (**p-r**) minocycline. For each subfigure, the results are represented with OD600 (left) and Bliss Interaction (right) heatmaps. Each heatmap represents the average of 3 replicates. For the Bliss Interaction Metric, positive values indicate synergy, negative values indicate antagonism, and neutral values indicate independence (additivism). * = interactions that are significantly different than additivism [p ≤ 0.05 by a one-sample non-equivalence t-test against the interval (−0.1, 0.1), with FDR-correction].

For antibiotics that target or disrupt the bacterial envelope – including polymyxin B (PXB), meropenem (MER), and ceftazidime (CAZ) – MCFAs consistently enhance antimicrobial activity. C6 and OCT each produced broad regions of positive Bliss scores when combined with PXB, indicating synergistic interactions across multiple stoichiometries (**Figure 6b,c**). Similar synergy zones were observed for CAZ and MER (**Figure 6e,f,h,i**), reflecting possible enhancement of cell wall or membrane stress when these agents were used together. These synergistic patterns were stoichiometrically-dependent but reproducible across replicates, supporting a cooperative interaction between MCFAs and envelope-active drugs.

In contrast, interactions between MCFAs and antibiotics that target intracellular processes were largely additive. For trimethoprim–sulfamethoxazole (SXT) and levofloxacin (LEV), MCFA co-treatment produced mostly neutral Bliss values, with occasional isolated positive or negative regions (**Figure 6k,l,n,o**), consistent with general independence between fatty acid stress and either folate biosynthesis or DNA replication inhibition.

The most striking pattern emerged with minocycline (MIN), an inhibitor of translation: both hexanoate and octanoate produced strong and widespread antagonistic interactions across a broad range of dosing combinations (**Figure 6q,r**). These negative Bliss values indicate that MCFA exposure diminishes minocycline efficacy, a phenomenon that may reflect shifts in membrane energetics, efflux behavior, or ribosomal engagement under fatty acid stress.

Together, these results reveal a consistent pattern: MCFAs synergize with membrane-active or envelope-targeting antibiotics, display largely additive interactions with agents that inhibit DNA or folate metabolism, and antagonize antibiotics that block protein synthesis. This pattern parallels the MCFA-phage interactions observed earlier and suggests that MCFA driven alterations of the bacterial envelope potentiate extracellularly acting antimicrobials while inhibiting the activity of ribosome-targeting drugs.

## DISCUSSION

Medium-chain fatty acids address several problems that currently limit phage therapy by modifying bacterial susceptibility to phage via multiple mechanisms. At concentrations that fall within ranges previously investigated for topical antimicrobial formulations and well below concentrations associated with many pharmaceutical formulas and food applications, MCFAs can potentiate phage activity across a diverse group of both Gram-negative and Gram-positive pathogens, including STM, *S. aureus*, *Enterococcus* spp., and *P. aeruginosa*. Working primarily with STM, we find that MCFAs inhibit biofilm formation, delay or prevent phage-resistant bacterial rebound over 48 hours, and synergize with some antibiotics. This work elevates MCFAs as potential therapeutic accompaniments that stabilize and broaden phage efficacy, and highlights the general potential of small molecules as bacterial-directed phage adjuvants.

The strongest effects were observed in *E. faecalis*, where 10 mM OCT combined with phage CAR yielded synergy across nearly all strains tested, including many that were recalcitrant to phage alone, effectively expanding the functional host range of CAR in our panel. Mechanistically, *E. faecalis* is unusually poised to exploit environmental fatty acids via a multi-component fatty acid kinase (FakAB) pathway, which promotes rapid uptake and incorporation of exogenous fatty acids into membrane phospholipids (Dong 2025; Zou 2024). This may alter membrane fluidity and surface architecture in ways that heighten susceptibility to envelope-acting stresses and facilitate phage adsorption/entry. Notably, synergy in *E. faecalis* emerged only at a low starting biomass, indicating that host-density-dependent factors (rather than MOI alone) govern MCFA-phage cooperation.

In contrast, MCFA-phage combinations in *S. aureus* exhibited a pronounced stoichiometric dependence (with zones of synergy, additivity, and antagonism) and in *P. aeruginosa* they were consistently additive. For *S. aureus,* low OCT with higher phage input favored synergy, whereas high OCT produced antagonism. This stoichiometric dependence mirrors reports for phage-antibiotic combinations in *S. aureus*, where synergy was observed between phages and envelope-active antibiotics at certain stoichiometries and antagonism at others (Kunz 2024), and is consistent with broader phage-antibiotic synergy studies showing that mechanism-of-action and stoichiometry jointly govern cooperation vs. interference (Liu 2020). One explanation is that moderate MCFA levels fluidize the membrane and subtly remodel the envelope, increasing exposure/accessibility of wall teichoic acid (WTA) – the canonical receptor for many staphylococcal phages (Xia 2011) and the putative receptor for phage Ludmila (Kumar, in prep) – easing adsorption/entry. Excess MCFA may over-perturb the envelope, trigger depolarization and leakage, and/or mask or structurally disrupt WTA, thereby reducing receptor integrity and impairing early phage steps (Parsons 2012). Whatever the reason behind this stoichiometric dependence, it suggests that systematic dose-response grids will be needed as part of any therapeutic development.

We also found that environmental context heavily impacts phage-MCFA interactions. In synthetic CF sputum medium (SCFM1), we observed that MCFAs are more bioactive, yet phage-MCFA interactions are largely additive across multiple STM-phage pairs (**Figs. 2**, **S2**, **4**), with notable phage-specific shifts (for example, SBP2Φ2 and KB824 are more effective while ANB28 and ESCSMPI are less effective relative to BHI). In the effort to mimic CF-like nutritional and physicochemical cues, growth in SCFM1 appears to reprogram envelope and community phenotypes in ways that both enhance MCFA killing and constrain additional synergy from phages. Nutrient cues in sputum-like conditions can reprogram pathogen physiology and thereby shift the phage-adjuvant interaction regime, emphasizing the importance of airway-relevant media when inferring translational potential (Neve 2021; Meneses 2025; Palmer 2007). Growth in SCFM1 may also change phage receptor landscapes and penetration constraints, possibly explaining why SBP2Φ2 outperforms while ANB28 underperforms in SCFM1 versus BHI.

Overall, our results indicate that MCFA-phage cooperativity is broad (although not universal) across taxa and the orderly chain-length trend observed in STM supports a general physicochemical mechanism rather than one driven by FAs targeting specific bacterial receptors. Several changes result from MCFA exposure, each of which may impact phage efficacy depending on the specific bacterial species/strain and the nutritional environment. Mechanistically, we focused our investigations on three separate (but interconnected) explanations: cell envelope remodeling, biofilm perturbation, and suppression of phage resistance.

Consistent with an envelope-focused mechanism, MCFAs showed strong synergy with cell-envelope-active antibiotics (**Fig 6**) – including polymyxin B (which disrupts lipopolysaccharide and cell membrane integrity) (Borrelli 2025) and ceftazidime (a third-generation cephalosporin antibiotic that inhibits cell-wall synthesis) (Hayes 1983). Borrelli et al. show that polymyxin B lethality requires remodeling of the outer membrane and loss of lipopolysaccharide (LPS), which enables subsequent inner-membrane permeabilization. Because MCFAs increase membrane disorder and permeability (Parsons 2012), they likely lower the threshold for this LPS-remodeling step, providing a biophysical explanation for the strong synergy we observe between MCFAs and polymyxin B. By contrast, we observed antagonism with the translation inhibitor minocycline, potentially reflecting FA-induced efflux that also exports minocycline (Rosenberg 2003). Together, these patterns indicate that MCFA synergy peaks with agents that exploit envelope destabilization (e.g., polymyxin B, ceftazidime), and this same logic explains their cooperation with phages: by lowering the biophysical threshold for outer-membrane and cell-wall failure, MCFAs increase receptor access, accelerate DNA delivery, and blunt envelope-based defenses.

Consistent with prior work showing that fatty acids disrupt biofilms by perturbing the extracellular matrix (Lin 2023, Kumar 2020, Nazzaro 2026) and that sub-lethal phage exposure can promote biofilm formation and enable persistence (Fernandez 2017, Burkle 2025, Hansen 2019), we find that MCFAs significantly reduce biofilm biomass and that this effect persists even with a biofilm-inducing phage (ANB28). This offers a second, biofilm-centric explanation to the observed cooperation: by thinning the extracellular matrix and altering surface physicochemistry, MCFAs may improve receptor exposure and mass transport, increasing productive virion-bacterial encounters that sustain phage predation over time. In line with this interpretation, biofilm architecture strongly modulates phage access (Winans 2022), underscoring the value of pre-emptively destabilizing the matrix. Similarly, other biofilm-targeting adjuncts (e.g., endolysins) reduce *P. aeruginosa* biofilms and are being explored as co-therapies (Nayab 2025), reinforcing the principle that matrix-weakening agents potentiate phage activity. Collectively, these observations indicate that by dismantling or preventing the formation of biofilms, MCFAs convert biofilm-protected communities into phage-permissive populations, thereby amplifying adsorption and sustaining predation.

In concert with enhanced adsorption and matrix access, these effects translate into a third outcome: MCFAs suppress the emergence of phage-resistant subpopulations across strains, phages, and media (**Fig. 2**). Where rebound typically reflects envelope-modifying mutations or biofilm-mediated refuge, MCFA co-treatment eliminates or narrows these escape routes. For example, STM strain k279a, which is partially refractory to ANB28, showed little suppression with phage alone; yet as little as 5 mM OCT markedly depressed growth, and higher concentrations prevented regrowth for 48 h. Likewise, in STM strain B28B and phage SBP2Φ2, the characteristic ∼12 h rebound was delayed or abolished by 15 mM OCT. Mechanistically, destabilizing the outer membrane, exposing receptors, and disrupting protective matrix architecture reduces the spatial and physiological refuges where resistance arises, while increasing the vulnerability of emerging mutants to ongoing phage attack. Together, these effects constrain the expansion of resistant subpopulations, transforming transient killing into sustained population control.

MCFAs emerge as practical phage co-agents that can broaden functional host range, delay or prevent resistance outgrowth, and increase phage efficacy under clinically relevant conditions. By testing multiple strains from four diverse pathogens we understand that phage cooperativity is broad but not universal, and that interactions vary in surprising ways, for example the stoichiometric dependence in *S. aureus*, and the host-density dependence in *E. faecalis*. By highlighting the promise of fatty acids specifically, our work argues for further research into (i) chemotype expansion beyond straight-chain saturated MCFAs to include unsaturation, branching, and headgroup engineering; and (ii) formulation science to overcome solubility and delivery limitations (e.g., micelles, liposomes, dry-powder inhalation, mucus-penetrating nanoparticles). Our attempts at systematically testing longer chain fatty acids (C11 and above) were hampered by poor solubility. Longer chain fatty acids might be even more effective as phage adjuvants than the MCFAs examined here, but this will require different chemistries or solubilization strategies. All work discussed here was performed *in vitro*, and while SCFM1 provides a physiologically relevant model of the cystic fibrosis airway environment, it does not fully recapitulate host factors such as immune activity, spatial structure, or mucus heterogeneity.

Together, our results show that MCFA-induced envelope and matrix remodeling can make phages work better, longer, and against more strains, establishing MCFAs as prototype molecular adjuvants for phage therapy. Fatty acids are widely used in foods and some medicines, however co-administration with phages (especially via airway delivery) requires dedicated safety, tolerability, and pharmacokinetic studies, including epithelial compatibility, aerosol performance, and mucociliary clearance. Beyond MCFAs, our findings support a general strategy of deploying molecular adjuvants to tune phage-host interactions. These advances suggest a tractable discovery space in which fatty acids, cations, bacterial metabolites, and pathway-specific inhibitors are rationally paired with phages to improve adsorption, entry, replication, and population-level outcomes.

## MATERIALS AND METHODS

### Bacterial Cultures

*Stenotrophomonas maltophilia, Staphylococcus aureus,* and *Pseudomonas aeruginosa* strains were stored at −80°C in 25% glycerol stocks and streaked onto fresh plates every 3–5 weeks during periods of active use. *Enterococcus faecalis,* and *Enterococcus faecium* were maintained as frozen stocks in 96-well plates (Dunham 2026, under review) at −80°C. For all experiments, with the exception of the host range experiments, bacterial cultures were inoculated from fresh plates into brain heart infusion BHI (CAT #RPI-B11000-1000.0) media and grown overnight at 37°C with shaking. Cultures were subsequently diluted 1:10 and grown to log phase prior to use in all assays except the host range assays. Optical density (OD_600_) measurements of liquid cultures in tubes were obtained using a Go Direct SpectroVis Plus Spectrophotometer before use in a plate.

### Phage Stocks

Phage stocks were stored in 10% glycerol at −80°C in 0.5 mL aliquots and thawed on ice immediately prior to use. Phage stocks were generated by using the plate propagation methods outlined in Bonilla et. al. (Bonilla 2016). In this method, high titer phage is added to a soft agar overlay with a permissive host to create web lysis plates. SM buffer (2-3 mL) is added to each plate, then the SM and top agar are scraped and collected into a tube that is spun down at 10,000 RPM for 10 minutes to separate bacteria from phage. The bacterial pellet is discarded and the supernatant containing the phage is pipetted off, syringe filtered through a 0.2 um filter, and aliquoted into 0.6 mL tubes. Glycerol is added to create final glycerol concentrations of 10% by volume and the phages are stored at −80°C. Titer data of the phage stocks was collected when freeze-thaw cycles began to impact phage performance and each time a new batch of phage was used.

### Fatty Acid Preparation

Fatty acids were purchased from Fisher Scientific, VWR and Sigma-Aldrich and used without further purification. Catalog numbers and sources for all compounds are provided in **Supplemental Table 3**. For experiments with pH-uncontrolled fatty acids, 1 M primary stock solutions were prepared directly in brain heart infusion (BHI) broth (Cat# RPI-B11000-1000.0). For experiments with pH-neutral fatty acids, 1 M primary stocks were prepared by combining each fatty acid with its conjugate base at ratios calculated using the Henderson–Hasselbalch equation to achieve a target pH near neutrality. Stocks were prepared in either BHI or Synthetic Cystic Fibrosis Sputum medium (SCFM1: SynthBiome Product # 10001-500; https://www.synthbiome.com/) The final pH was verified to be between 7.2 - 7.5 using pH indicator strips (Cat# 14850121). Fatty acid stocks prepared in BHI were stored at room temperature in a dark, enclosed environment, while stocks prepared in SCFM1 were stored at 4°C wrapped in aluminum foil to minimize light exposure.

### Checkerboard Synergy Assays

Checkerboard assays were performed in 96-well, non-treated, clear flat-bottom plates (Fisher Scientific Cat# 0877253) to assess interactions between phages and fatty acids (**Figure 1a**). Phage stocks were serially diluted in SM buffer and 10uL of phage dilutions were added down columns of the plate, with the highest concentration added across row A, lowest concentration added across row F, and an SM buffer control (plus FA) across row G. 6 media controls (200 μL BHI) and bacteria only controls (110 μL BHI and 90 μL bacteria) were added to row H. Log-phase bacterial cultures were diluted in BHI to an OD_600_ of 0.185 +/− 0.005. 90 μL of the diluted bacterial culture was added to each experimental well. The 1 M primary stock solution of pH neutral fatty acid was added to BHI to make working solutions. These working solutions were twice the desired final concentration in the well, as we added 100 μL of these dilutions to each well to create a total volume of 200uL. The lowest concentration of fatty acid was in column 2, the highest concentration in column 12, and a BHI-vehicle control in column 1. Plates were sealed with BreathEasy membranes (Cat # NC0674973) and incubated at 37°C in humidified containers with a beaker of DI water to prevent desiccation. OD_600_ measurements were collected after 24 hours +/− 1 hour with a Cerillo Alto plate reader (Cerillo Cat# ALT1600).

Analysis of checkerboard synergy data was performed in R (via RStudio 2023.09.1). For the data in **Figure 1**, .xlsx files containing the raw OD_600_ values were imported using readxl (Wickham 2019), reshaped via melt() from the reshape2 package (Wickham 2007), and joined and ordered via dplyr functions (Wickham 2023). For each compound, the media-only wells were taken from Row H, Columns 7–12. Outliers were removed using the IQR rule (1.5×IQR), with the remaining values defining the media reference used for hypothesis testing. To compute the OD growth statistics, for each condition, we tested whether the mean of its three replicates exceeded the media reference distribution using one-sided Welch two-sample *t*-tests with alternative “greater”; *p*-values were recorded and non-significant conditions (indicating a well that was not significantly different than 0; *p* > 0.05) labeled with “Θ” in **Figure 1**. Welch’s unequal-variance *t* is appropriate when the focal and reference samples differ in variance and size (Welch 1947).

Background-corrected OD values (***OD_bkg_***) were calculated by subtracting the outlier-robust mean of media-only wells (Row H, Columns 7-12) from every well:

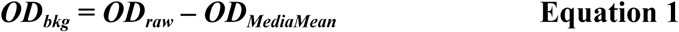

For each experimental well, the observed survival fraction (***S_obs_***), which was calculated as the ratio of the experimental ***OD_bkg_*** to the corresponding bacteria-only ***OD_bkg_*** from well G1:

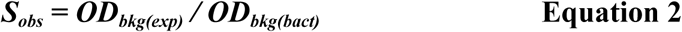

Following the original Bliss formulation (Bliss 1939) and modern treatments of statistical synergy (Demidenko 2019), the expected surviving fraction (***S_exp_***) for a two-agent combination is defined as the product of the observed single-agent survival fractions at the same doses, thus accounting for the individual antimicrobial effects of each agent:

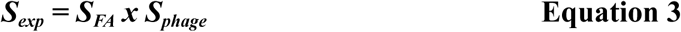

And here we define the Bliss Interaction Metric (***Bliss***) as the difference between the ***S_exp_*** and ***S_obs_***:

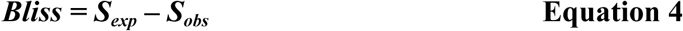

By this sign convention, positive ***Bliss*** values indicate synergy (observed survival less than independent expectation), zero indicates additivity, and negative indicates antagonism (Demidenko 2019). To flag combinations that depart meaningfully from additivity, we used an interval-null approach: for each plate position, we tested whether the replicate mean ***Bliss*** lay outside [-0.1 – + 0.1] units via two one-sided *t*-tests (evidence for µ < −0.1 or µ > 0.1); the smaller *p*-value was reported, and cells with *p* ≤ 0.05 were labeled with “**\***”. This follows the logic of two one-sided testing for equivalence/non-equivalence (adapted here to detect non-equivalence to additivity) and uses Welch’s *t* under unequal variances (Lakens 2017).

Where relevant we also calculated percent reduction in growth (***%R***) relative to bacteria-only growth as:

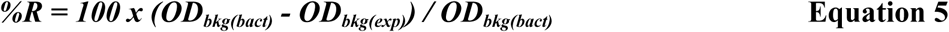

In summaries of single-agent concentration series, we compared each dose to 0 mM by two-sample *t*-tests and adjusted the family of *p*-values using the Holm sequentially-rejective method (as implemented in p.adjust(method=“holm”)) (Holm 1979).

We rendered heatmaps using ggplot2, with diverging palettes and fixed limits for the ***Bliss*** scale. For ***Bliss***, we centered the palette at 0 and padded the color scale so that the interval ±0.1 maps to a neutral tone, visually marking additivity. Panel assembly and figure export used ggpubr and scales (for color/label utilities). For figure-ready panels, we employed minimalist themes and, where needed, ggtext (element_markdown) to style facet labels.

### Dual Checkerboard Synergy Assays

Two types of dual-checkerboard synergy assays were performed to compare interaction landscapes across media or treatment conditions within a single plate. In each assay, the OCT concentrations were held constant for both halves, while the condition of interest differed between the left and right halves of the plate (e.g., BHI vs. SCFM1 for identical *S. maltophilia*–phage pairings). pH-neutral OCT 1M primary stock solutions were prepared in the corresponding media and added across columns at final concentrations of 0, 1, 2.5, 5, 10, and 15 mM. Phages were serially diluted 10-fold across rows, with the highest titer added to the top row (the highest titer varied by phage: 10⁹ PFU/mL for ANB28 and 10⁵–10⁸ PFU/mL for SBP2Φ2 depending on the host strain).

Following a similar structure, dual-condition checkerboard assays were also performed to test pH-neutral fatty acid-antibiotic interactions. A single acid was used per plate (acetic acid [C2], hexanoic acid [C6], or octanoic acid [OCT]) and was applied across the x-axis at final concentrations of 0, 1, 2.5, 5, 10, and 20 mM. Two antibiotics were tested per plate, with one antibiotic assigned to each half (left vs. right), including ceftazidime/minocycline, meropenem/levofloxacin, and polymyxin B/trimethoprim–sulfamethoxazole pairings.

Antibiotics were obtained from Thomas Scientific, Neta Scientific, Sigma-Aldrich, and Fisher Scientific (see **Supplemental Table 3**). Meropenem, levofloxacin, and minocycline were reconstituted in DMSO, while trimethoprim–sulfamethoxazole was provided as a liquid stock in DMSO. Polymyxin B was supplied in aqueous solution and filter-sterilized prior to use. All antibiotic stocks were stored at −20°C and diluted in 0.9% saline such that addition of 10 μL per well yielded the final concentrations indicated on the y-axis of each assay.

Analysis of the dual checkerboard synergy assay analysis was performed via methods analogous to those described above for the checkerboard synergy assays.

### Growth Curves

Growth curve assays were performed in clear flat-bottom 96-well plates using the same bacterial preparation described above. Log-phase cultures were diluted to an OD_600_ of 0.185 +/− 0.005 and 90 µL of the bacterial suspension was added to each well. Row A contained control conditions, including bacteria-only (90μL bacteria, 10μL SM buffer, 100μL BHI) wells on the left and bacteria+phage (90μL bacteria, 10μL phage, and 100μL BHI) wells on the right. For the remaining rows (B–H), pH-neutral fatty acids were added across the whole plate in increasing concentrations down the plate to achieve final concentrations of 0, 2.5, 5, 10, 12.5, and 15 mM (the 2.5 and 12.5 mM conditions were omitted from figures and analysis for clarity). Phage treatment (10μL phage of interest) was applied to the right half of the plate, while the left half contained the no-phage control (10 μL SM buffer). Growth curves were conducted for two strain–phage pairs (*S. maltophilia* B28B with phage SBP2Φ2 and *S. maltophilia* K279a with phage ANB28) in both BHI and artificial sputum medium SCFM1, resulting in four experimental plates. Plates were incubated at 37°C in a Cerillo Alto plate reader, and OD₆₀₀ measurements were collected every 10 minutes for 48 hours.

Growth curves were plotted in R using ggplot2, with LOESS-smoothed overlays where appropriate, and assembled with ggpubr (Kassambara 2020). The growthcurver (Sprouffske 2020) package was used to generate growth curve metrics.

### Biofilm Assays

Biofilm assays were performed using a static crystal violet staining assay based on a protocol from O’Toole & Kolter’s original microtiter/tube-style method (O’Toole & Kolter 1998). Strain K279a was grown overnight in brain heart infusion (BHI) broth at 37°C with aeration, diluted into fresh BHI the following day, and grown to logarithmic phase before normalization to an OD₆₀₀ of 0.100 ± 0.05. For each biological replicate, 2 mL of standardized culture was distributed into sterile capped culture tubes under four conditions: bacteria only (20 µL SM buffer + 20 µL BHI), bacteria + phage (20 µL ANB28 phage, 10⁵ PFU per tube + 20 µL BHI), bacteria + octanoic acid (20 µL SM buffer + 20 µL 1 M octanoic acid to achieve a final concentration of 10 mM), and combination treatment (20 µL phage + 20 µL octanoic acid). When phage or octanoic acid were not included in the treatment, 20 µL of the appropriate SM buffer or BHI were used instead. Media controls of 980 µL BHI and 20 µL SM were included for background subtraction and baseline measurements.

Tubes were incubated statically at 37°C for 48 hours, only being removed once at 24 hours for OD₆₀₀ measurements. Following incubation, OD₆₀₀ was read using the Vernier spectrophotometer in the tubes they were incubated in (48 Hour measurement). Then, without disrupting wall-adhered biofilms, all liquid was carefully transferred via pipette to a new tube, homogenized, and OD₆₀₀ values were read (Total Biomass measurement). 2.5 mL of 1% crystal violet in Ultra Pure Water was added to the original tubes to stain the wall-adhered biofilm over the course of 10 minutes. The stain was carefully poured out and the tubes were rinsed in 2 water baths until the liquid from the tube ran clear. The tubes were tapped upside down to remove excess water and left upside down to dry overnight. Once dry, 3 mLs of 30% acetic acid was added to all tubes, mixed 3 times, and left for 10 minutes. After calibrating with 30% acetic acid, the OD_590_ values were read and recorded (Attached Biofilm measurement).

Biofilm assay data were analyzed in R using the tidyverse framework. Measurements were compared across conditions at each time point using pairwise Wilcoxon rank-sum tests with Benjamini–Hochberg false discovery rate (FDR) method. Data were visualized using boxplots with overlaid jittered points to display individual replicates.

### Host Range Analysis

Overnight cultures of host range panels were created by inoculating 5 μL of glycerol stock from freezer template plates into 600 μL of BHI in deep-well plates, which were sealed with a breathable non-woven rayon seal (Fisher Scientific #1256705) and incubated overnight at 37°C with shaking. The following day, cultures were diluted with 700 uL of fresh BHI and used to inoculate experimental 96-well plates in triplicate. Each plate was divided into four vertical sections representing bacteria-only, bacteria + phage, bacteria + octanoic acid (OCT), and combination treatments. Wells contained 90 μL bacterial culture supplemented with either SM buffer or phage and either BHI or pH-neutral OCT working solutions to achieve the desired conditions. Plates also included media controls in rows A and H. Plates were sealed with BreathEasy membranes (Cat# NC0674973), incubated statically at 37°C in humidified containers, and OD₆₀₀ was measured at ∼24 h.

Sub-inhibitory concentrations of phage and OCT were determined empirically for each species to enable detection of cooperative effects. Final conditions included 10 mM OCT with 10⁵ PFU/mL phage CAR for *Enterococcus faecalis*, 20 mM OCT with 10³ PFU/mL phage Ludmila for *Staphylococcus aureus*, and 15 mM OCT with 10⁵ PFU/mL phage PAPS1 for *Pseudomonas aeruginosa*. Once host range data was collected and analyzed, strains of interest were selected for closer examination through checkerboard assays as they allow for a range of phage and OCT dilutions. Notably, for *E. faecalis*, assays were performed using a lower initial inoculum (approximately 1:10,000 dilution of overnight cultures) to match parallel experimental workflows. When strains were selected for further checkerboard assays, synergistic patterns did not match host range data unless initial inocula was diluted 1:10,000.

Analysis for the host range assays was performed as described for the checkerboard synergy assays, but with the following differences. For each replicate, background subtraction to generate the OD_bkg_ values were computed by using the outlier-robust media mean (IQR, 1.5 × IQR rule) of 5 media control wells. For the STM assays in **Figure 4** strains were ordered by phylogenetic relatedness, and for all other host range assays (e.g., those in **Figures 5** and **S5**) they were ordered by the mean OD_600_ in the combined condition. Statistics were computed per strain across replicates in the phage & FA condition. For %R, we tested whether the mean differed from 0 using one-sample t-tests, applying Benjamini–Hochberg FDR correction across strains and marking FDR < 0.05 with “*”. For Bliss, we used an interval-null of [−0.1,+0.1] via two one-sided t-tests(evidence for μ < −0.1 or μ > +0.1); the smaller p-value was BH-adjusted across strains and significant strains (FDR < 0.05) were labeled with “*” in the Bliss heatmap. Plots were generated with ggplot2, panel assembly with ggpubr, and utilities from readxl, reshape2, dplyr, and scales (Wickham 2020).

## ACKNOWLEDGEMENTS

We thank Cameron Miller, Georgiana Kourepenos, Jason Rothman, and all past and present members of the Whiteson lab who contributed to this project. We also thank Heather Maughan for thoughtful editorial feedback. K.R.R. was supported by the Department of Education’s training grant Graduate Assistance in the Areas of National Need (GAANN) fellowship (P200A240034) and the National Institutes of Health training grant T32GM152338. S.J.B.D was supported by the Cystic Fibrosis Foundation (003135F221) and by the Cystic Fibrosis Research Institute (PG13391). Additional research funding was provided by the Cystic Fibrosis Research Institute under award 5680618.

## Notes

### Competing Interest Statement

The authors have declared no competing interest.

## WORKS CITED

Antimicrobial Resistance Collaborators. “Global Burden of Bacterial Antimicrobial Resistance in 2019: A Systematic Analysis.” The Lancet, vol. 399, no. 10325, 12 Feb. 2022, pp. 629–655, www.thelancet.com/journals/lancet/article/PIIS0140-6736(21)02724-0/fulltext, 10.1016/S0140-6736(21)02724-0.

Bliss, C. I. “THE TOXICITY of POISONS APPLIED JOINTLY1.” Annals of Applied Biology, vol. 26, no. 3, Aug. 1939, pp. 585–615, 10.1111/j.1744-7348.1939.tb06990.x.

Barrio-Pujante, Antonio, et al. “Auxin (Indole-3-Acetic Acid) Modulation of Quorum Sensing Enhances Phage Susceptibility in Klebsiella Pneumoniae.” European Journal of Clinical Microbiology & Infectious Diseases, 27 Mar. 2026, 10.1007/s10096-026-05454-z. Accessed 3 Apr. 2026.

Bonilla, Natasha, et al. “Phage on Tap–a Quick and Efficient Protocol for the Preparation of Bacteriophage Laboratory Stocks.” PeerJ, vol. 4, 26 July 2016, www.ncbi.nlm.nih.gov/pmc/articles/PMC4975003/, 10.7717/peerj.2261.

Borrelli, Carolina, et al. “Polymyxin B Lethality Requires Energy-Dependent Outer Membrane Disruption.” Nature Microbiology, 29 Sept. 2025, 10.1038/s41564-025-02133-1.

Bürkle, Magdalena, et al. “Phage-Phage Competition and Biofilms Affect Interactions between Two Virulent Bacteriophages and Pseudomonas Aeruginosa.” The ISME Journal, vol. 19, no. 1, June 2025, p. wraf065, pubmed.ncbi.nlm.nih.gov/40188480/, 10.1093/ismejo/wraf065.

Clokie, Martha R.J., et al. “Phages in Nature.” Bacteriophage, vol. 1, no. 1, Jan. 2011, pp. 31–45, pmc.ncbi.nlm.nih.gov/articles/PMC3109452/, 10.4161/bact.1.1.14942.

Desbois, Andrew P., and Valerie J. Smith. “Antibacterial Free Fatty Acids: Activities, Mechanisms of Action and Biotechnological Potential.” Applied Microbiology and Biotechnology, vol. 85, no. 6, 3 Dec. 2009, pp. 1629–1642, link.springer.com/article/10.1007%2Fs00253-009-2355-3, 10.1007/s00253-009-2355-3.

Demidenko, Eugene, and Todd W. Miller. “Statistical Determination of Synergy Based on Bliss Definition of Drugs Independence.” PLOS ONE, vol. 14, no. 11, 25 Nov. 2019, p. e0224137, 10.1371/journal.pone.0224137. Accessed 3 Dec. 2019.

Dhanda, Geetika, et al. “Antibiotic Adjuvants: A Versatile Approach to Combat Antibiotic Resistance.” ACS Omega, vol. 8, no. 12, 14 Mar. 2023, 10.1021/acsomega.3c00312.

Dong, Huijuan, et al. “Defining the *Enterococcus Faecalis* Fatty Acid Kinase System of Exogeneous Fatty Acid Utilization.” Molecular Microbiology, 2 Aug. 2025, 10.1111/mmi.70017. Accessed 8 Sept. 2025.

Dunham SJB, Willkeen GA, Darby B, Corley JM, Hahn A, Klapper I, Bean HD, Caverly LJ, Thornton CS, Martin C, Quinn RA, Widder S, Bailey BA, Wagner BD, Garg N, Planet PJ, Hunter RC, LiPuma JJ, Rohwer F, Whiteson KL. 2026. The Guild Model of CF Airway Microbial Ecology. mBio17:e03668–25.10.1128/mbio.03668-25

Elshazly A, Russell DA, Hatfull GF, Schooley RT, Bollyky PL, Pourtois JD, Freeman KG. 2025 Conference on Bacteriophages: Biology, Dynamics, and Therapeutics. Pathog Immun. 2026 Feb 9;11(1):69–95. doi: 10.20411/pai.v11i1.942. PMID: 41704570; PMCID: PMC12908700.

Eskenazi, Anaïs, et al. “Combination of Pre-Adapted Bacteriophage Therapy and Antibiotics for Treatment of Fracture-Related Infection due to Pandrug-Resistant Klebsiella Pneumoniae.” Nature Communications, vol. 13, no. 1, 18 Jan. 2022, p. 302, www.nature.com/articles/s41467-021-27656-z, 10.1038/s41467-021-27656-z.

Fernández, Lucía, et al. “Low-Level Predation by Lytic Phage PhiIPLA-RODI Promotes Biofilm Formation and Triggers the Stringent Response in Staphylococcus Aureus.” Scientific Reports, vol. 7, no. 1, 19 Jan. 2017, 10.1038/srep40965. Accessed 29 Mar. 2022.

Fletcher, Joshua R., et al. “Commensal-Derived Short-Chain Fatty Acids Disrupt Lipid Membrane Homeostasis in *Staphylococcus Aureus*.” MBio, vol. 17, no. 1, 14 Jan. 2026, 10.1128/mbio.01392-25. Accessed 24 Mar. 2026.

Gallagher, Tara, et al. “Cystic Fibrosis-Associated Stenotrophomonas Maltophilia Strain-Specific Adaptations and Responses to PH.” Journal of Bacteriology, vol. 201, no. 7, Apr. 2019, 10.1128/jb.00478-18. Accessed 11 Mar. 2025.

Ghorbani, Peyman, et al. “Short-Chain Fatty Acids Affect Cystic Fibrosis Airway Inflammation and Bacterial Growth.” European Respiratory Journal, vol. 46, no. 4, 28 May 2015, pp. 1033–1045, 10.1183/09031936.00143614. Accessed 20 May 2021.

Hansen, Mads Frederik, et al. “Big Impact of the Tiny: Bacteriophage–Bacteria Interactions in Biofilms.” Trends in Microbiology, vol. 27, no. 9, Sept. 2019, pp. 739–752, www.cell.com/trends/microbiology/fulltext/S0966-842X(19)30099-X, 10.1016/j.tim.2019.04.006. Accessed 25 Jan. 2020.

Hayes, M. V., and D. C. Orr. “Mode of Action of Ceftazidime: Affinity for the Penicillin-Binding Proteins of Escherichia Coli K12, Pseudomonas Aeruginosa and Staphylococcus Aureus.” The Journal of Antimicrobial Chemotherapy, vol. 12, no. 2, 1 Aug. 1983, pp. 119–126, pubmed.ncbi.nlm.nih.gov/6413485/, 10.1093/jac/12.2.119. Accessed 12 Oct. 2021.

Holm, Sture. “A Simple Sequentially Rejective Multiple Test Procedure.” Scandinavian Journal of Statistics, vol. 6, no. 2, 1979, pp. 65–70, www.jstor.org/stable/4615733.

Kassambara, Alboukadel. “Ggpubr: “Ggplot2” Based Publication Ready Plots.” R-Packages, 13 Feb. 2020, cran.r-project.org/web/packages/ggpubr/index.html.

Koskella, Britt, and Sean Meaden. “Understanding Bacteriophage Specificity in Natural Microbial Communities.” Viruses, vol. 5, no. 3, 11 Mar. 2013, pp. 806–823, 10.3390/v5030806.

Kotlyarov, Stanislav. “Role of Short-Chain Fatty Acids Produced by Gut Microbiota in Innate Lung Immunity and Pathogenesis of the Heterogeneous Course of Chronic Obstructive Pulmonary Disease.” International Journal of Molecular Sciences, vol. 23, no. 9, 26 Apr. 2022, p. 4768, 10.3390/ijms23094768.

Kumar, Prasun, et al. “Fatty Acids as Antibiofilm and Antivirulence Agents.” Trends in Microbiology, vol. 28, no. 9, Sept. 2020, pp. 753–768, 10.1016/j.tim.2020.03.014.

Kumar, Ritwik, et al. “Phage Therapy from Bench-to-Bedside: The Characterization of Staphylococcus phage OCPT-Ludmila and its use to treat MRSA Sinusitis: In Prep. 2026

Kunz Coyne, Ashlan J., et al. “Exploring Synergistic and Antagonistic Interactions in Phage-Antibiotic Combinations against ESKAPE Pathogens.” Microbiology Spectrum, vol. 12, no. 10, 3 Oct. 2024, pmc.ncbi.nlm.nih.gov/articles/PMC11468199/pdf/spectrum.00427-24.pdf, 10.1128/spectrum.00427-24.

Lin, Wen-Chun, et al. “Octanoic Acid Promotes Clearance of Antibiotic-Tolerant Cells and Eradicates Biofilms of Staphylococcus Aureus Isolated from Recurrent Bovine Mastitis.” Biofilm, vol. 6, 12 Aug. 2023, pp. 100149–100149, 10.1016/j.bioflm.2023.100149. Accessed 11 June 2025.

Lakens, Daniël. “Equivalence Tests: A Practical Primer for T Tests, Correlations, and Meta-Analyses.”Social Psychological and Personality Science, vol. 8, no. 4, May 2017, pp. 355–362, 10.1177/1948550617697177.

Liu, Carmen Gu, et al. “Phage-Antibiotic Synergy Is Driven by a Unique Combination of Antibacterial Mechanism of Action and Stoichiometry.” MBio, vol. 11, no. 4, 25 Aug. 2020, 10.1128/mBio.01462-20.

Ma, Li, et al. “Metals Enhance the Killing of Bacteria by Bacteriophage in Human Blood.” Scientific Reports, vol. 8, no. 1, 2 Feb. 2018, www.ncbi.nlm.nih.gov/pmc/articles/PMC5797145/pdf/41598_2018_Article_20698.pdf, 10.1038/s41598-018-20698-2.

McCutcheon, Jaclyn G., and Jonathan J. Dennis. “The Potential of Phage Therapy against the Emerging Opportunistic Pathogen Stenotrophomonas Maltophilia.” Viruses, vol. 13, no. 6, 3 June 2021, p. 1057, 10.3390/v13061057.

Meneses, Luciana, et al. “Directed Evolution of Phages in Biofilms Enhances Pseudomonas Aeruginosa Control through Improved Lipopolysaccharide Recognition.” Nature Communications, vol. 16, no. 1, 20 Nov. 2025, pp. 10219–10219, www.nature.com/articles/s41467-025-65014-5, 10.1038/s41467-025-65014-5.

Monsibais, Alisha N, et al. “Enhanced Suppression of Stenotrophomonas Maltophilia by a Three-Phage Cocktail: Genomic Insights and Kinetic Profiling.” Antimicrobial Agents and Chemotherapy, vol. 69, no. 3, 22 Jan. 2025, www.researchgate.net/publication/388276833_Enhanced_suppression_of_Stenotrophomonas_m altophilia_by_a_three-phage_cocktail_genomic_insights_and_kinetic_profiling, 10.1128/aac.01162-24.

Moon, Kyung, et al. “Considerations and Perspectives on Phage Therapy from the Transatlantic Taskforce on Antimicrobial Resistance.” Nature Communications, vol. 16, no. 1, 4 Dec. 2025, pp. 10883–10883, 10.1038/s41467-025-64608-3.

Naghavi, M., Vollset, S. E., Ikuta, K. S., Swetschinski, L. R., Gray, A. P., Wool, E. E., Robles Aguilar, G., Mestrovic, T., Smith, G., Han, C., Hsu, R. L., Chalek, J., Araki, D. T., Chung, E., Raggi, C., Gershberg Hayoon, A., Davis Weaver, N., Lindstedt, P. A., Smith, A. E., … Murray, C. J. L. (2024). Global burden of bacterial antimicrobial resistance 1990–2021: A systematic analysis with forecasts to 2050. The Lancet, 404(10459), 1199–1226. 10.1016/s0140-6736(24)01867-1

Nayab, Sehrish, et al. “Synergism of Phages and Antimicrobial Peptides for Treating Multidrug Resistant Bacterial Pathogens.” Exploration of Drug Science, vol. 3, 12 Nov. 2025, 10.37349/eds.2025.1008133. Accessed 16 Nov. 2025.

Nazzaro, Filomena, et al. “Fatty Acids as Prebiotics and Their Role in Antibiofilm Activity.” Antibiotics, vol. 15, no. 1, 5 Jan. 2026, p. 57, 10.3390/antibiotics15010057. Accessed 8 Jan. 2026.

Neve, Rachel L., et al. “Impact of Artificial Sputum Medium Formulation on Pseudomonas Aeruginosa Secondary Metabolite Production.” Journal of Bacteriology, vol. 203, no. 21, 12 Oct. 2021, 10.1128/jb.00250-21. Accessed 17 Nov. 2022.

O’Toole, George A., and Roberto Kolter. “Initiation of Biofilm Formation InPseudomonas FluorescensWCS365 Proceeds via Multiple, Convergent Signalling Pathways: A Genetic Analysis.” Molecular Microbiology, vol. 28, no. 3, Apr. 1998, pp. 449–461, onlinelibrary.wiley.com/doi/pdf/10.1046/j.1365-2958.1998.00797.x, 10.1046/j.1365-2958.1998.00797.x.

Palmer, Kelli L., et al. “Nutritional Cues Control Pseudomonas Aeruginosa Multicellular Behavior in Cystic Fibrosis Sputum.” Journal of Bacteriology, vol. 189, no. 22, 14 Sept. 2007, pp. 8079–8087, 10.1128/jb.01138-07.

Parsons, J. B., et al. “Membrane Disruption by Antimicrobial Fatty Acids Releases Low-Molecular-Weight Proteins from Staphylococcus Aureus.” Journal of Bacteriology, vol. 194, no. 19, 27 July 2012, pp. 5294–5304, dx.doi.org/10.1128%2FJB.00743-12, 10.1128/jb.00743-12.

Perikleous, Evanthia P., et al. “Antibiotic Resistance in Patients with Cystic Fibrosis: Past, Present, and Future.” Antibiotics, vol. 12, no. 2, 20 Jan. 2023, p. 217, www.ncbi.nlm.nih.gov/pmc/articles/PMC9951886/, 10.3390/antibiotics12020217.

Pirnay, Jean-Paul, et al. “Personalized Bacteriophage Therapy Outcomes for 100 Consecutive Cases: A Multicentre, Multinational, Retrospective Observational Study.” Nature Microbiology, vol. 9, no. 6, 1 June 2024, pp. 1434–1453, www.nature.com/articles/s41564-024-01705-x, 10.1038/s41564-024-01705-x.

Pompilio, Arianna, et al. “Phenotypic and Genotypic Characterization of Stenotrophomonas Maltophilia Isolates from Patients with Cystic Fibrosis: Genome Diversity, Biofilm Formation, and Virulence.” BMC Microbiology, vol. 11, no. 1, 5 July 2011, 10.1186/1471-2180-11-159. Accessed 15 Mar. 2022.

Reasoner, Seth A., et al. “Urinary Tract Infections in Cystic Fibrosis Patients.” Journal of Cystic Fibrosis, vol. 21, no. 1, 1 Jan. 2022, pp. e1–e4, www.sciencedirect.com/science/article/pii/S1569199321013035, 10.1016/j.jcf.2021.07.005. Accessed 25 May 2022.

Rosenberg, Emiko Y., et al. “Bile Salts and Fatty Acids Induce the Expression of Escherichia Coli AcrAB Multidrug Efflux Pump through Their Interaction with Rob Regulatory Protein.” Molecular Microbiology, vol. 48, no. 6, 15 May 2003, pp. 1609–1619, 10.1046/j.1365-2958.2003.03531.x. Accessed 19 Dec. 2020.

Rountree, P. M. “The Role of Divalent Cations in the Multiplication of Staphylococcal Bacteriophages.” Journal of General Microbiology, vol. 12, no. 2, 1 Apr. 1955, pp. 275–287, 10.1099/00221287-12-2-275. Accessed 4 Feb. 2023.

Ryan, Robert P., et al. “The Versatility and Adaptation of Bacteria from the Genus Stenotrophomonas.” Nature Reviews Microbiology, vol. 7, no. 7, July 2009, pp. 514–525, 10.1038/nrmicro2163.

Sprouffske, Kathleen. “Simple Metrics to Summarize Growth Curves [R Package Growthcurver Version 0.3.1].” R-Project.org, 19 Oct. 2020, cran.r-project.org/web/packages/growthcurver/index.html, https://cran.r-project.org/package=growthcurver. Accessed 10 Apr. 2026.

Terlizzi, Vito, et al. “Stenotrophomonas Maltophilia in People with Cystic Fibrosis: A Systematic Review of Prevalence, Risk Factors and Management.” European Journal of Clinical Microbiology & Infectious Diseases : Official Publication of the European Society of Clinical Microbiology, vol. 42, no. 11, Nov. 2023, pp. 1285–1296, pubmed.ncbi.nlm.nih.gov/37728793/, 10.1007/s10096-023-04648-z.

Uyttebroek, Saartje, et al. “Safety and Efficacy of Phage Therapy in Difficult-To-Treat Infections: A Systematic Review.” The Lancet Infectious Diseases, vol. 0, no. 0, 3 Mar. 2022, www.thelancet.com/journals/laninf/article/PIIS1473-3099(21)00612-5/fulltext#articleInformatio n, 10.1016/S1473-3099(21)00612-5.

Van Nieuwenhuyse, Brieuc, et al. “Bacteriophage-Antibiotic Combination Therapy against Extensively Drug-Resistant Pseudomonas Aeruginosa Infection to Allow Liver Transplantation in a Toddler.” Nature Communications, vol. 13, no. 1, 29 Sept. 2022, p. 5725, www.nature.com/articles/s41467-022-33294-w, 10.1038/s41467-022-33294-w.

Verma, Anjali, et al. “Short-Chain Fatty Acid (SCFA) as a Connecting Link between Microbiota and Gut-Lung Axis─a Potential Therapeutic Intervention to Improve Lung Health.” ACS Omega, vol. 9, no. 13, 19 Mar. 2024, pp. 14648–14671, 10.1021/acsomega.3c05846.

Waters, Valerie, et al. “Stenotrophomonas Maltophiliain Cystic Fibrosis.” American Journal of Respiratory and Critical Care Medicine, vol. 183, no. 5, Mar. 2011, pp. 635–640, 10.1164/rccm.201009-1392oc.

Welch, B. L. “The Generalization of “Student’s” Problem When Several Different Population Variances Are Involved.” Biometrika, vol. 34, no. 1-2, 1947, pp. 28–35, 10.1093/biomet/34.1-2.28.

Wickham H, Bryan J (2025). readxl: Read Excel Files. R package version 1.4.5, https://github.com/tidyverse/readxl, https://readxl.tidyverse.org.

Wickham, Hadley. “Reshaping Data with ThereshapePackage.” Journal of Statistical Software, vol. 21, no. 12, 2007, 10.18637/jss.v021.i12.

Wickham, Hadley, et al. “Dplyr: A Grammar of Data Manipulation.” R-Packages, 18 Aug. 2020, cran.r-project.org/web/packages/dplyr/index.html.

Wickham, Hadley, et al., “Scales: Scale Functions for Visualization.” R-Packages April 2025, https://cran.r-project.org/web/packages/scales/index.html

Winans, James B., et al. “Multispecies Biofilm Architecture Determines Bacterial Exposure to Phages.” PLOS Biology, vol. 20, no. 12, 22 Dec. 2022, p. e3001913, 10.1371/journal.pbio.3001913.

Wong, Joel W H, and Emily P Balskus. “Small Molecules as Modulators of Phage-Bacteria Interactions.” Current Opinion in Chemical Biology, vol. 84, Feb. 2025, p. 102566, pubmed.ncbi.nlm.nih.gov/39736196/, 10.1016/j.cbpa.2024.102566.

Xia, G., et al. “Wall Teichoic Acid-Dependent Adsorption of Staphylococcal Siphovirus and Myovirus.” Journal of Bacteriology, vol. 193, no. 15, 3 June 2011, pp. 4006–4009, 10.1128/jb.01412-10.

Zang, Zhiyu, et al. “Chemical Inhibition of a Bacterial Immune System.” Cell Host & Microbe, vol. 34, no. 2, 1 Jan. 2026, 10.1016/j.chom.2026.01.003. Accessed 2 Feb. 2026.

Zhang, Hongchao, et al. “Antimicrobial Action of Octanoic Acid against Escherichia Coli O157:H7 during Washing of Baby Spinach and Grape Tomatoes.” Food Research International, vol. 125, Nov. 2019, p. 108523, 10.1016/j.foodres.2019.108523. Accessed 13 Sept. 2021.

Zou, Qi, et al. “The Enteric Bacterium *Enterococcus Faecalis* Elongates and Incorporates Exogenous Short and Medium Chain Fatty Acids into Membrane Lipids.” Molecular Microbiology, vol. 122, no. 5, 8 Oct. 2024, pp. 757–771, 10.1111/mmi.15322. Accessed 16 Dec. 2024.

