## Supplemental Figures 1-5 for "Fatty acids synergize with bacteriophages against multidrug-resistant pathogens"

**Supplemental Information**

Figure S1. Growth inhibition of *S. maltophilia* strain K279a by SCFAs & MCFAs.
Figure S2. MCFAs rescue activity of phage ANB28 at low MOIs.
Figure S3. pH-neutral MCFAs inhibit STM growth in synthetic CF sputum media (SCFM1).
Figure S4. MCFAs are bactericidal and do not significantly impact phage viability.
Figure S5. MCFAs display predominantly additive and strain-dependent interactions with bacteriophages in *P. aeruginosa*.

a)

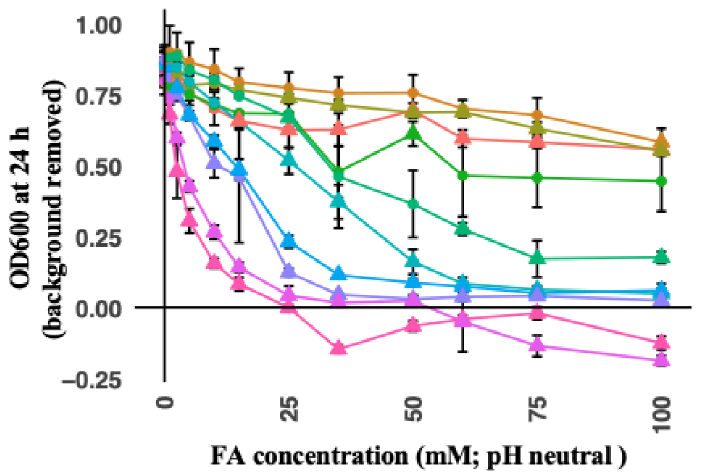


| | 50% GR [mM] | 95% GR [mM} |
|---|---|---|
| C1 | - | - |
| C2 | - | - |
| C3 | - | - |
| C4 | 65 | - |
| C5 | 50 | - |
| C6 | 35 | 65 |
| C7 | 15 | 65 |
| C8 | 10 | 35 |
| C9 | 2.5 | 25 |
| C10 | 2.5 | 25 |

b)

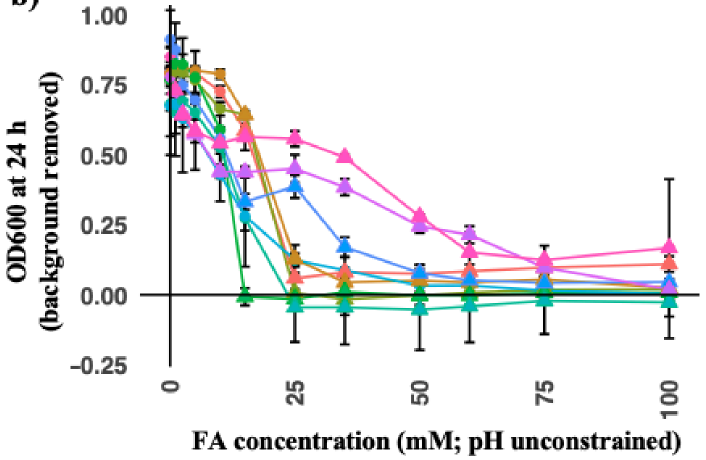


| | 50% GR [mM] | 95% GR [mM} | FA conc. at which pH ≤ 6 |
|---|---|---|---|
| C1 | 25 | 25 | 25 |
| C2 | 25 | 35 | 25 |
| C3 | 25 | 25 | 25 |
| C4 | 15 | 15 | 25 |
| C5 | 15 | 25 | 25 |
| C6 | 15 | 25 | 65 |
| C7 | 15 | 65 | 100 |
| C8 | 35 | 100 | >100 |
| C9 | 50 | - | >100 |

c)

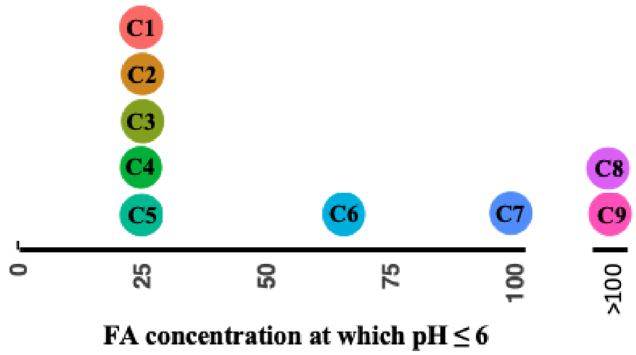


**Figure S1. Growth inhibition of *S. maltophilia* strain K279a by SCFAs & MCFAs.** (**a**) pH-neutral FAs; (**b**) acidic FAs. Shown are 24 hr $OD_{600}$ values with the background removed. The error bars show the standard deviation of 3 replicates for each condition; (**c**) dot plot depicting the FA concentration at which the pH is at or below 6. C10 could not be effectively solubilized in its acidic form. pH was tested using pH indicator strips. Each dot is labeled with FA chain length.

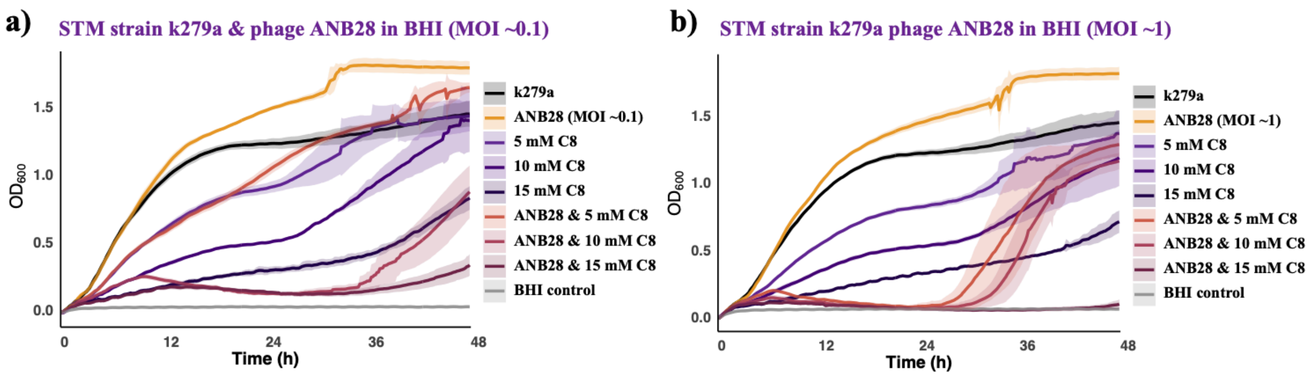


**Figure S2**. **MCFAs rescue activity of phage ANB28 at low MOIs**. 48 h growth curves for STM strain K279a with ANB28 at an approximate MOI of (**a**) 0.1 and (**b)** 1. All assays were conducted in BHI. Shown are the averages (solid lines) and standard deviations (shading) for 5 replicates. Phages were added at an approximate titer of 1 x $10^6$ pfu/mL for **a** and 1 x $10^7$ pfu/mL for **b**.

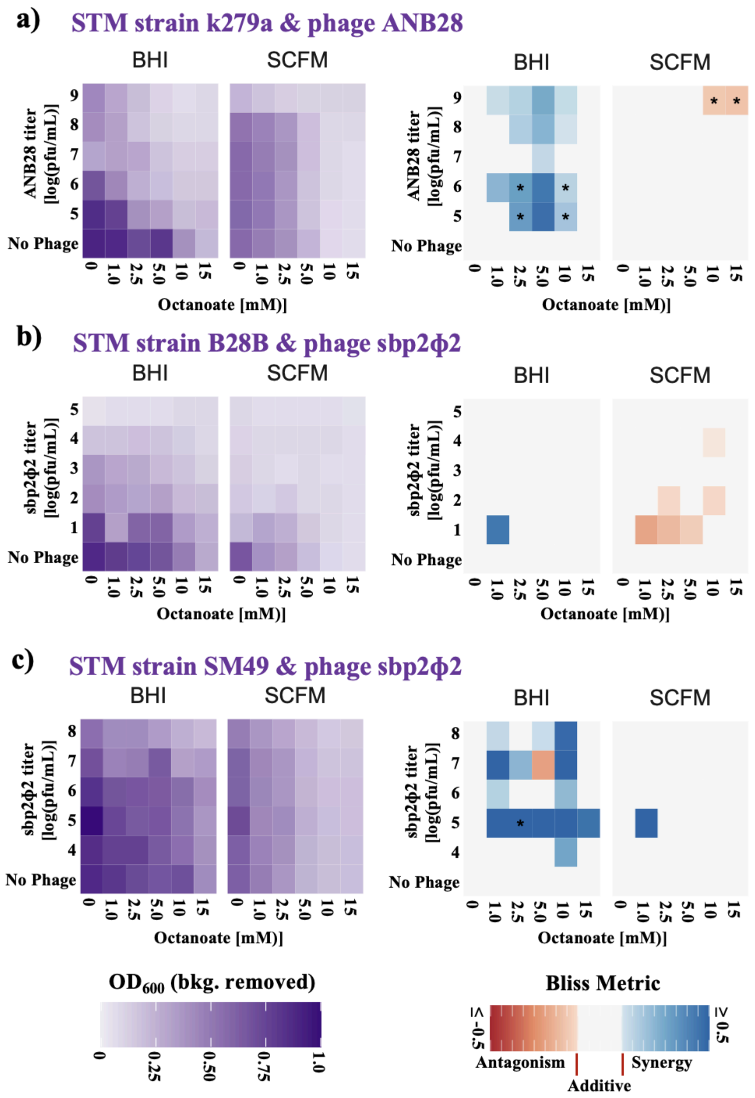


**Figure S3. pH-neutral MCFAs inhibit STM growth in synthetic CF sputum media (SCFM1)**. Checkerboard synergy assays between pH-neutral OCT and: (**a**) phage ANB28 against strain k279a, (**b**) phage SBP2ϕ2 against strain B28B, and (**c**) phage SBP2ϕ2 against strain SM49. Data represents the average of three replicates. A phage titer of 1 x $10^7$ pfu/mL corresponds to an approximate MOI of 1. For the Bliss Interaction Metric, positive values indicate synergy, negative values indicate antagonism, and neutral values indicate independence (additivism). Interactions that are significantly different than additivism [$p \leq 0.05$ by a one-sample non-equivalence t-test against the interval $(-0.1, 0.1)$] are indicated by “*” in the Bliss Interaction heatmaps.

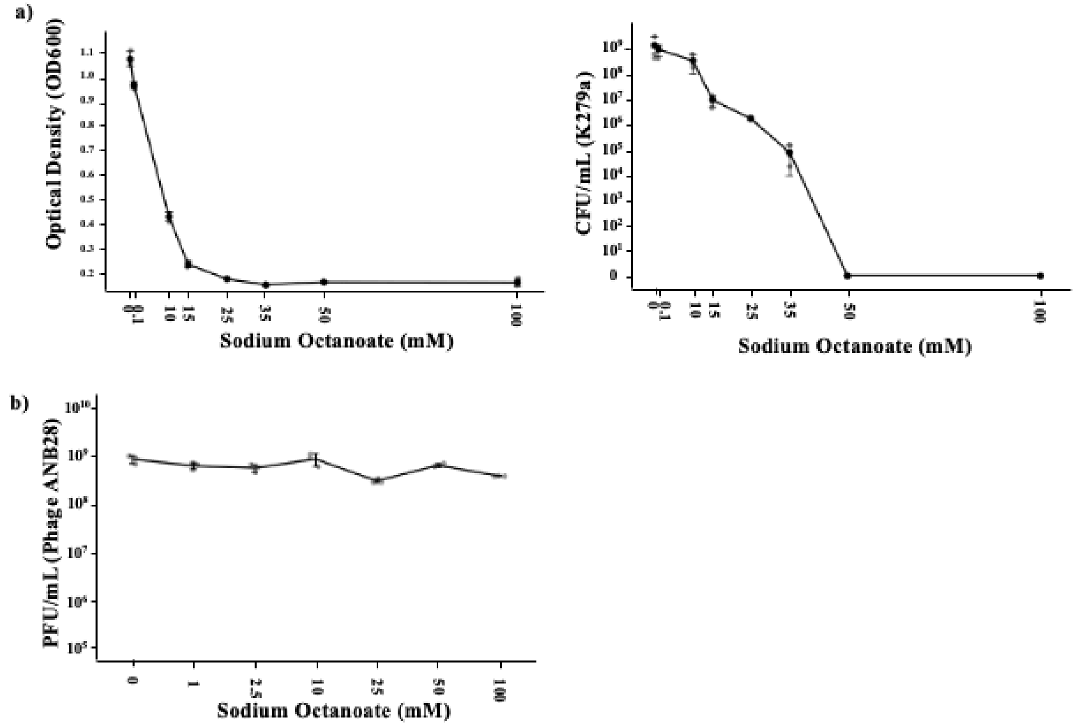


**Figure S4**. **MCFAs are bactericidal and do not significantly impact phage viability**. (**a**) $OD_{600}$ after 24 h incubation with varying concentrations of pH-neutral OCT (left) and subsequent colony counting (right). Strain K279a was incubated for 24 hours with varying concentrations of OCT, washed to remove residual OCT, and plated for colony counting. (**b**) Phage viability. Phage ANB28 was stored in an SM buffer with different concentrations of OCT at 4C. After two hours, the contents of each tube was further diluted in the SM buffer until reaching a $10^{-4}$ or $10^{-5}$ dilution. A soft agar overlay was performed with log phase k279a and incubated at 37℃ for 24 hours. The plaques were counted and used to calculate the titer of the phage in the tube.

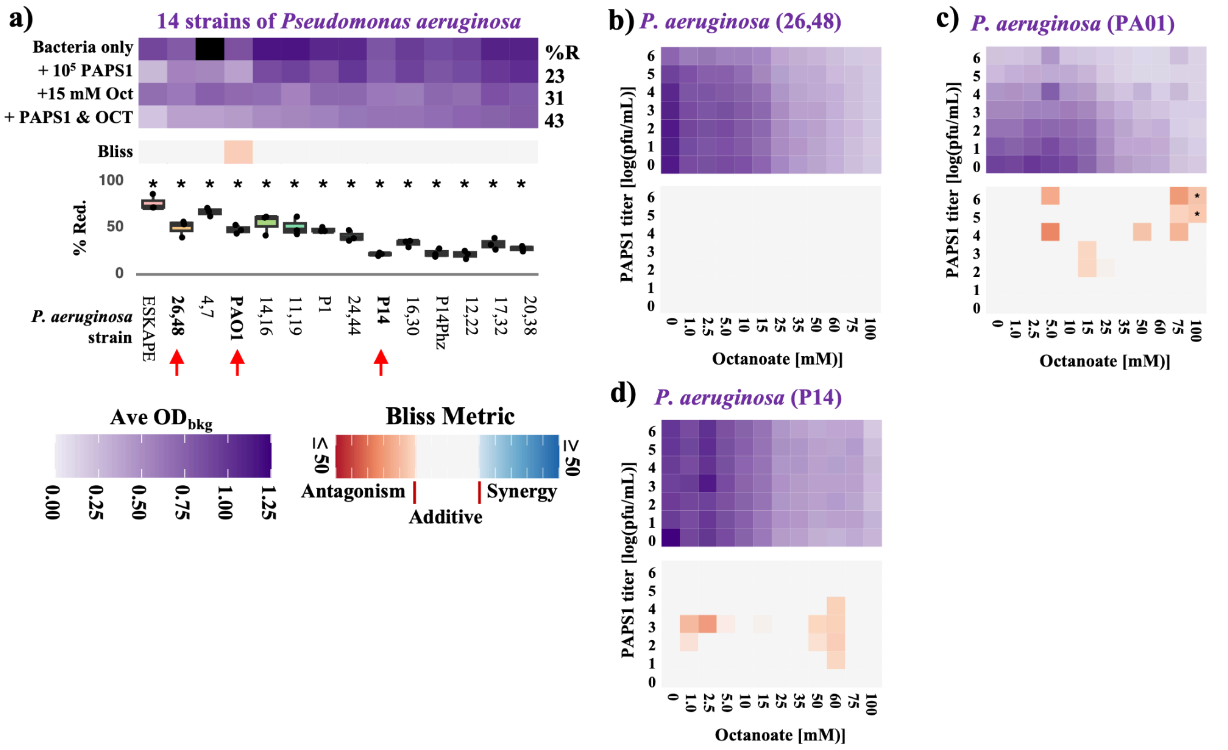


**Figure S5. MCFAs display predominantly additive and strain-dependent interactions with bacteriophages in *P. aeruginosa*.** (**a**) Multi-strain assay showing 24 h $OD_{600}$ measurements for 14 *P. aeruginosa* strains treated with phage PAPS1 ($10^5$ PFU/mL), OCT (15 mM), or the combination. Bliss interaction scores are shown below, with * indicating strains with interactions significantly different from additivity [$p \leq 0.05$ by a one-sample non-equivalence t-test against the interval (−0.1, 0.1), with FDR-correction]. (**b–d**) Checkerboard synergy assays for three representative *P. aeruginosa* strains (26,48; PAO1; and P14), showing $OD_{600}$ heatmaps (top) and corresponding Bliss Interaction Metrics (bottom). All data represent the mean of three replicates.
