## Supplemental Tables 1-3 for "Fatty acids synergize with bacteriophages against multidrug-resistant pathogens"

**Supplemental Information**

Supplemental Table 1: List of strains.
Supplemental Table 2: List of phages.
Supplemental Table 3: List of key reagents.

**Supplemental Table 1**: List of strains.

| Species | Strain | Source | Isolated from: | abx resistance profile |
|---|---|---|---|---|
| *Stenotrophomonas maltophilia* | K279a | STM type Strain | NCBI #ASM7248v1 | unknown |
| *Stenotrophomonas maltophilia* | B28B | UCSD | blood infection | unknown |
| *Stenotrophomonas maltophilia* | FL10 | UCI | Florida, Pulmonary infection | unknown |
| *Stenotrophomonas maltophilia* | Lipton23 | UCI | Pulmonary infection | unknown |
| *Stenotrophomonas maltophilia* | HLP | UCI | Pulmonary infection | unknown |
| *Stenotrophomonas maltophilia* | SM10KA | UCSD | Lesio MTAE8 CXBS | unknown |
| *Stenotrophomonas maltophilia* | SM12LS | UCSD | UR MTAE8 CXURN | unknown |
| *Stenotrophomonas maltophilia* | SM15KA | UCSD | BRONW MTAE8 CXQBR | unknown |
| *Stenotrophomonas maltophilia* | SM16LS | UCSD | SITE MAC-S CXBS | unknown |
| *Stenotrophomonas maltophilia* | SM20TB | UCSD | UR MACU CXURN | ceftazadime, levofloxacin |
| *Stenotrophomonas maltophilia* | SM25LS | UCSD | UR MAC-S CXURN | unknown |
| *Stenotrophomonas maltophilia* | SM26KA | UCSD | BRONW MAC CXRES | unknown |
| *Stenotrophomonas maltophilia* | SM27KA | UCSD | UR MTAE8 CXURN | ceftazadime |
| *Stenotrophomonas maltophilia* | SM32KA | UCSD | BRONW CNC-8 CXAFB | unknown |
| *Stenotrophomonas maltophilia* | SM49LS | UCSD | RESCF MTAE8 CXRES | levofloxacin |
| *Stenotrophomonas maltophilia* | SM50JS | UCSD | Rescf MTAE8 CXRES | resistant to all |
| *Stenotrophomonas maltophilia* | SM73PII | UCSD | CXRES | ceftazadime, levofloxacin |
| *Stenotrophomonas maltophilia* | SM74PII | UCSD | UR MAC-S CXURN | unknown |
| *Staphylococcus aureus* | 10,6 | UCI - Whiteson Lab Wound Collection | Left thumb (abscess) | unknown |
| *Staphylococcus aureus* | 12,21 | UCI - Whiteson Lab Wound Collection | Right femur | unknown |
| *Staphylococcus aureus* | 2,4 | UCI - Whiteson Lab Wound Collection | Left chin (abscess) | unknown |
| *Staphylococcus aureus* | 27,51 | UCI - Whiteson Lab Wound Collection | Sinus | unknown |

| | | | | |
|---|---|---|---|---|
| *Staphylococcus aureus* | 30,59 | UCI - Whiteson Lab Wound Collection | Scalp skin | unknown |
| *Staphylococcus aureus* | 32,63 | UCI - Whiteson Lab Wound Collection | Right ankle (drainage) | unknown |
| *Staphylococcus aureus* | 19,36 | UCI - Whiteson Lab Wound Collection | Left hand (skin lesion) | unknown |
| *Staphylococcus aureus* | 27,52 | UCI - Whiteson Lab Wound Collection | Sinus | unknown |
| *Staphylococcus aureus* | 9,15 | UCI - Whiteson Lab Wound Collection | Right eyebrow (abscess) | unknown |
| *Staphylococcus aureus* | 10,18 | UCI - Whiteson Lab Wound Collection | Left thumb (abscess) | unknown |
| *Staphylococcus aureus* | 21,40 | UCI - Whiteson Lab Wound Collection | Lower left leg (drainage) | unknown |
| *Staphylococcus aureus* | 25,46 | UCI - Whiteson Lab Wound Collection | Right arm (skin; drainage) | unknown |
| *Staphylococcus aureus* | 7,12 | UCI - Whiteson Lab Wound Collection | Abdomen (wound) | unknown |
| *Staphylococcus aureus* | JVS2311 | UCI Medical Center, Donald Forthal | MRSA; external medical device | unknown |
| *Staphylococcus aureus* | G565 ATCC 33591 | Greg Burkeen, FLR, SDSU | ? | unknown |
| *Staphylococcus aureus* | 31,60 | UCI - Whiteson Lab Wound Collection | Lower left leg (lesion) | unknown |
| *Staphylococcus aureus* | PedCF | Greg Burkeen, FLR, SDSU | | unknown |
| *Staphylococcus aureus* | USA300 | UCI - Whiteson Lab Wound Collection | ATCC BAA-1680 | unknown |
| *Pseudomonas aeruginosa* | 4,7 | UCI - Whiteson Lab Wound Collection | Sinus (chronic sinusitis) | unknown |
| *Pseudomonas aeruginosa* | 12,22 | UCI - Whiteson Lab Wound Collection | Right femur | unknown |
| *Pseudomonas aeruginosa* | 14,26 | UCI - Whiteson Lab Wound Collection | Left ear | unknown |
| *Pseudomonas aeruginosa* | 16,30 | UCI - Whiteson Lab Wound Collection | Sinus | unknown |
| *Pseudomonas aeruginosa* | 20,38 | UCI - Whiteson Lab Wound Collection | Nose (wound) | unknown |
| *Pseudomonas aeruginosa* | 24,44 | UCI - Whiteson Lab Wound Collection | Sinus | unknown |
| *Pseudomonas aeruginosa* | 26,48 | UCI - Whiteson Lab Wound Collection | Left thigh | unknown |
| *Pseudomonas aeruginosa* | PAnmFLR01 (P1) | KW (CF isolate UCSD 2014) | unknown | unknown |
| *Pseudomonas aeruginosa* | WT P14 | Siryaporn (O'Toole) | unknown | unknown |

| | | | | |
|---|---|---|---|---|
| *Pseudomonas aeruginosa* | P14 DPhz | Hochbaum / Newman | unknown | unknown |
| *Pseudomonas aeruginosa* | 11,19 | UCI - Whiteson Lab Wound Collection | Sputum | unknown |
| *Pseudomonas aeruginosa* | 17,32 | UCI - Whiteson Lab Wound Collection | Sinus | unknown |
| *Pseudomonas aeruginosa* | ESKAPE Pa | ESKAPE Pathogens | unknown | unknown |
| *Pseudomonas aeruginosa* | PAO1 | Dorrestein/Phelan | unknown | unknown |
| *Enterococcus faecalis* | EF01PII | David Pride | bone left foot | none |
| *Enterococcus faecalis* | EF02PII | David Pride | urine | none |
| *Enterococcus faecalis* | EF03PII | David Pride | urine | gentamicin-syn |
| *Enterococcus faecium* | EF04PII | David Pride | blood | amp, penicillin G |
| *Enterococcus faecium* | EF05PII | David Pride | urine | amp, penicillin G, vancomycin |
| *Enterococcus faecium* | EF06PII | David Pride | blood picc line | amp, penicillin G, tetracycline, vanco |
| *Enterococcus faecalis* | EF07PII | David Pride | urine | gentamicin-syn |
| *Enterococcus faecalis* | EF08PII | David Pride | blood | none |
| *Enterococcus faecalis* | EF09PII | David Pride | urine | tetracycline, vanco |
| *Enterococcus faecalis* | EF10PII | David Pride | body site lesion leg | none |
| *Enterococcus faecium* | EF11PII | David Pride | body site pancreatic fluid | linezolid |
| *Enterococcus faecalis* | EF12PII | David Pride | bile duct fluid | none |
| *Enterococcus faecalis* | EF13PII | David Pride | body site abscess | none |
| *Enterococcus faecalis* | EF14PII | David Pride | urine | gentamicin-syn |
| *Enterococcus faecalis* | EF15PII | David Pride | urine | none |
| *Enterococcus faecium* | EF16PII | David Pride | urine | none |
| *Enterococcus faecalis* | EF17PII | David Pride | urine | none |
| *Enterococcus faecium* | EF18PII | David Pride | body site pancreas | amp, penicillin G, tetracycline, vanco |
| *Enterococcus faecalis* | EF19PII | David Pride | urine | none |
| *Enterococcus faecium* | EF20PII | David Pride | body fluid wound | amp, penicillin G, tetracycline, vanco |
| *Enterococcus faecalis* | EF23PII | David Pride | urine | gentamicin-syn |
| *Enterococcus faecalis* | EF24PII | David Pride | abscess | none |
| *Enterococcus faecalis* | EF25PII | David Pride | blood | none |
| *Enterococcus faecalis* | EF28PII | David Pride | urine | none |
| *Enterococcus faecalis* | EF29PII | David Pride | urine | none |
| *Enterococcus faecalis* | EF30PII | David Pride | urine | none |
| *Enterococcus faecalis* | EF31PII | David Pride | urine | none |

| | | | | |
|---|---|---|---|---|
| *Enterococcus faecalis* | EF32PII | David Pride | urine | none |
| *Enterococcus faecalis* | EF33PII | David Pride | urine | none |
| *Enterococcus faecalis* | EF35PII | David Pride | urine | none |
| *Enterococcus faecalis* | EF36PII | David Pride | urine | none |
| *Enterococcus faecalis* | EF37PII | David Pride | urine | gentamicin-syn |
| *Enterococcus faecalis* | EF38PII | David Pride | urine | gentamicin-syn |
| *Enterococcus faecalis* | EF39PII | David Pride | bile duct fluid | none |
| *Enterococcus faecalis* | EF40PII | David Pride | urine | none |
| *Enterococcus faecium* | EF41PII | David Pride | body site swab | amp, gentamicin-syn, penicillin G, tetra, vanco |
| *Enterococcus faecalis* | EF42PII | David Pride | body site abdomen | gentamicin-syn |
| *Enterococcus faecalis* | EF43PII | David Pride | urine | none |
| *Enterococcus faecalis* | EF44PII | David Pride | urine | none |
| *Enterococcus faecalis* | EF45PII | David Pride | urine | none |
| *Enterococcus faecalis* | EF46PII | David Pride | urine | none |
| *Enterococcus faecalis* | EF47PII | David Pride | urine | gentamicin-syn |
| *Enterococcus faecium* | EF48PII | David Pride | urine | amp, penicillin G, tetracycline, vanco |
| *Enterococcus faecalis* | EF49PII | David Pride | urine | gentamicin-syn |
| *Enterococcus faecium* | EF50PII | David Pride | urine | amp, penicillin G, vanco |
| *Enterococcus faecalis* | EF51PII | David Pride | urine | none |
| *Enterococcus faecalis* | EF52PII | David Pride | urine | none |
| *Enterococcus faecium* | EF53PI | David Pride | aspirate abdominal fluid | none |
| *Enterococcus faecalis* | EF54PII | David Pride | aspirate abdominal fluid | none |
| *Enterococcus faecalis* | EF55PII | David Pride | urine | none |
| *Enterococcus faecalis* | EF57PII | David Pride | drainage ear | none |
| *Enterococcus faecalis* | EF58PII | David Pride | aortic abscess | gentamicin-syn |
| *Enterococcus faecalis* | EF59PII | David Pride | urine | none |
| *Enterococcus faecalis* | EF61PII | David Pride | urine | gentamicin-syn |
| *Enterococcus faecium* | EF62PII | David Pride | body tissue ulcer | amp, penicillin G, tetra, vanco |
| *Enterococcus faecalis* | EF63PII | David Pride | body site mouth swab | unknown |
| *Enterococcus faecalis* | EF64PII | David Pride | urine | none |
| *Enterococcus faecalis* | EF65PII | David Pride | urine | none |
| *Enterococcus faecalis* | EF66PII | David Pride | urine | none |
| *Enterococcus faecalis* | EF67PII | David Pride | urine | gentamicin-syn |
| *Enterococcus faecalis* | EF68PII | David Pride | urine | none |
| *Enterococcus faecalis* | EF69PII | David Pride | urine | gentamicin-syn |
| *Enterococcus faecalis* | EF70PII | David Pride | urine | gentamicin-syn |

| | | | | |
|---|---|---|---|---|
| *Enterococcus faecalis* | EF71PII | David Pride | urine | none |
| *Enterococcus faecalis* | EF72PII | David Pride | urine | none |
| *Enterococcus faecalis* | EF73PII | David Pride | urine | none |
| *Enterococcus faecalis* | EF74PII | David Pride | peritoneal fluid | minocycline, gentamycin-syn |
| *Enterococcus faecalis* | EF75PII | David Pride | aspirate abdominal fluid | none |
| *Enterococcus faecalis* | EF76PII | David Pride | aspirate abdominal fluid | none |
| *Enterococcus faecalis* | EF77PII | David Pride | urine | none |
| *Enterococcus faecalis* | EF78PII | David Pride | unknown | unknown |
| *Enterococcus faecium* | EF79PII | David Pride | body site swab | amp, penicillin G, tetra, vanco |
| *Enterococcus faecalis* | EF80PII | David Pride | unknown | unknown |
| *Enterococcus faecalis* | EF81PII | David Pride | body tissue finer | none |
| *Enterococcus faecalis* | EF82PII | David Pride | urine | none |
| *Enterococcus faecalis* | EF83PII | David Pride | urine | none |
| *Enterococcus faecalis* | EF84PII | David Pride | urine | gentamycin-syn |
| *Enterococcus faecium* | EF85PII | David Pride | blood | amp, penicllin G |
| *Enterococcus faecalis* | EF86PII | David Pride | urine | none |
| *Enterococcus faecalis* | EF87PII | David Pride | urine | none |
| *Enterococcus faecalis* | EF88PII | David Pride | urine | none |
| *Enterococcus faecium* | DP17 | David Pride | unknown | unknown |
| *Enterococcus faecalis* | Yi6 | BEI | unknown | unknown |
| *Enterococcus faecium* | TX1330 | BEI | unknown | unknown |
| *Enterococcus faecalis* | V587 | BEI | unknown | unknown |
| *Enterococcus faecalis* | DP05 | David Pride | unknown | unknown |
| *Enterococcus faecalis* | DP12 | David Pride | unknown | unknown |
| *Enterococcus faecalis* | DP15 | David Pride | unknown | unknown |
| *Enterococcus faecalis* | EF102PII | David Pride | unknown | unknown |
| *Enterococcus faecalis* | EF116PII | David Pride | unknown | unknown |
| *Enterococcus faecalis* | ESKAPE E. faecalis | ESKAPE Pathogens | unknown | unknown |

**Supplemental Table 2**: List of phages.

| Host organism | Phage name | Host strain | Isolation Source | Isolated By | Morphology | Genome Ref | Citation |
|---|---|---|---|---|---|---|---|
| *S. maltophilia* | ANB28 | B28B | Escondido Wastewater | Pride Lab-Alisha Blanc | Podovirus | PQ434692 | Monsibais 2025 |
| *S. maltophilia* | KB824 | B28B | Escondido Wastewater | Pride Lab | Siphovirus | PQ434693 | Monsibais 2025 |
| *S. maltophilia* | SPB2Φ2 | B28B | Escondido Wastewater | Pride Lab | Podovirus | PQ074228 | Monsibais 2025 |
| *S. maltophilia* | ESCSMPI | SM71PII | Escondido Wastewater | OCPT- Coby Fan | Siphovirus | | |
| *E. faecalis* | CAR | Yi6 | Escondido Wastewater | OCPT-Stephen Wandro | Myovirus | | |
| *S. aureus* | Ludmila | USA300 | SCCWRP wastewater | OCPT-Alexandra Albert | Podovirus | | |
| *P. aeruginosa* | PAPS1 | ESKAPE PA | Escondido Wastewater | OCPT-Ritwik Kumar | Unknown | | |

**Supplemental Table 3**: List of key reagents.

| **Type** | **Supplier** | **Compound** | **Product number** |
|---|---|---|---|
| Acid | Sigma | Formic acid | 695076-100ML |
| Acid | Fluka | Acetic acid | 49199-50ML-F |
| Acid | Fisher | Propionic acid | S25508 |
| Acid | VWR | Butyric acid | TCB0754-25ML (EA) |
| Acid | VWR | Pentanoic acid | TCV0003-25ML (EA) |
| Acid | VWR | Hexanoic acid | TCH0105-25ML (EA) |
| Acid | VWR | Heptanoic acid | TCH0030-25ML (EA) |
| Acid | VWR | Octanoic acid | TCO0027-25ML (EA) |
| Acid | VWR | Nonanoic acid | TCP0952-25ML (EA) |
| Acid | Fisher | Decanoic acid | D001725G |
| Base | Sigma | Sodium formate | 247596-100G |
| Base | Sigma | Sodium acetate | S2889-250G |
| Base | Sigma | Sodium propionate | P1880-100G |
| Base | Sigma | Sodium butyrate | 303410-5G |
| Base | Ambeed, Inc. | Sodium pentanoate | AMBH2D6FA579-25G |
| Base | Sigma | Sodium hexanoate | C4026-10G |
| Base | Ambeed, Inc. | Sodium heptanoate | AMBH9614F4B7-25G |
| Base | Sigma | Sodium octanoate | C5038-10G |
| Base | Sant Cruz Biotech | Sodium nonanoate | 296399 |
| Base | Sigma | Sodium decanoate | C4151-5G |
| Antibiotic | Thomas Scientific | Meropenam | C817F47 (EA/1) |
| Antibiotic | Sigma-Aldrich | Cotrimoxazole (SXT) | A2487-1ML |
| Antibiotic | Thomas Scientific | Levofloxacin | C790J97 |
| Antibiotic | Thomas Scientific | Minocyclin | C828G93 |
| Antibiotic | Neta Scientific | Ceftazidime Hydrate | RPI-C59200-1.0 |
| Antibiotic | Fisher | Polymixin B Sulfate | AAJ671638EQ |
